# Tick hazard in the South Downs National Park (UK): species, distribution, key locations for future interventions, site density, habitats

**DOI:** 10.1101/2021.03.23.436533

**Authors:** Jo Middleton, Ian Cooper, Anja S. Rott

## Abstract

**Background:** The South Downs National Park (SDNP) is the UK’s most visited National Park, and a foci of tick-borne Lyme disease. A range of human pathogens have been detected in UK ticks and related hosts, and the first presumed autochthonous cases of tick-borne encephalitis and babesiosis were recorded in 2019–20. SDNP’s key objectives include conserving wildlife and encouraging enjoyment of the countryside, so interventions are needed that reduce hazard without negatively affecting ecosystem health. To be successful these require knowledge of site hazards, and we aimed to provide this to enable action.

**Methods:** British Deer Society volunteers submitted ticks removed from deer. Key potential intervention sites were selected and ticks collected by drag-sampling six 50 m^2^ transects per site, in most cases twice yearly for two years. Ticks were identified in-lab (sex, life stage, species), hazard measured as tick presence, Density of Ticks (all life stages, DOT), and Density of Nymphs (DON). Sites and habitat types were analysed for association with hazard. Distribution across SDNP was mapped in a Geographic Information System (GIS), by combining and comparing our fieldwork results with records from five other data sources (recent and historic).

**Results:** 87 *Ixodes ricinus* (all but one adults, 82%F) were removed from 14 deer (*Dama dama* n=10; *Capreolus capreolus* n=3; 1 not recorded; tick burden, 1–35) at 12 locations (commonly woodland). Five potential key intervention sites were identified and drag-sampled 2015–16, collecting 623 ticks (238 on-transects): 53.8% nymphs, 42.5% larvae, 3.7% adults (13M, 10F). Ticks were present on-transects at all sites drag-sampled *(I.ricinus* at three, *Haemaphysalis punctata* at two). The Mens (TM, the quietist site for human visitors) had the highest DOT at 30/300 m^2^ (DON=30/300 m^2^), followed by Queen Elizabeth Country Park (QECP, the busiest) at 22/300 m^2^ (12/300 m^2^), Cowdray Estate (CE) at 8/300 m^2^ (6/300 m^2^), and Seven Sisters Country Park (SSCP) at 1/300 m^2^ (1/300 m^2^). Ditchling Beacon Nature Reserve (DBNR) was sampled 2016 only (one adult *H.punctata* collected). Woodland had significantly higher hazard than grazed downland, but ticks were present at all downland sites drag-sampled. GIS mapping showed *I.ricinus* identified in 33/37 of SDNPs 10 km^2^ grid squares, *Ixodes hexagonus* 10/37, *H.punctata* 7/37, *Dermacentor reticulatus* 1/37.

**Conclusions:** Mapping shows tick hazard is broadly distributed across SDNP. *Ixodes ricinus*was most common, though the seeming range expansion of *H.punctata* is concerning, particularly as it seems to thrive better on grazed downland than *I.ricinus*. Site specific recommendations include: management of small high hazard plots with heavy visitor numbers (QECP); signage on post-visit precautions (all sites); repellent impregnated clothing for deerstalkers (CE); flock trials to control *H.punctata* (SSCP, DBNR). Further research at TM, which has high tick density, may contribute to knowledge on ecological dynamics underlying infection density, and the potential use of predator re-introduction/protection as a public health intervention. Ecological research on *H.punctata* would aid control. The SDNP Authority is ideally placed to link and champion site-based and regional policies to reduce hazard, whilst avoiding or reducing conflict between public health and ecosystem health.

## INTRODUCTION

The South Downs National Park (SDNP) covers 1,627 km^2^ of the south-east of the British Isles, across Hampshire, West Sussex, and East Sussex. It encompasses two bioregions, the 140 km long chalk ridge of the South Downs, and the wooded lowland Weald. Though much of the Park is subject to industrial agriculture, substantive fragments of rare and species rich semi-natural chalk grassland can be found on its windy hills, whilst some of its woodlands are truly ancient (>1000 y) and harbor biodiverse ecological communities (Crane & Williams, 2013). It is the most visited national park in the UK, with an estimated 39 million visitor days per year (NPUK, 2014), *c.*120,000 people live and/or work within its borders, two million live within 5 km (SDNPA, 2020). A substantial part of the national park is private land with limited or no public access (Bangs, 2008). However, the area is crisscrossed by 3218 km of public rights-of-way (TTC, 2018), and there is sizable local authority owned country parks and myriad nature reserves. Some stretches of downland are legally classed as ‘Access Land’ (SDS, 2021) and some landowners also allow permissive paths. The South Downs Way is one of 15 UK national trails and is very popular with walkers, cyclists, and horse-riders. Over one year 61,191 people were counted passing one point of the trail (ESCC, 2016), locations closer to carparks can be far busier still (HCC, 2020).

Ticks (Ixodida) are second only to mosquitoes globally as vectors of human pathogens (Lawrie *et al*., 2004). Twenty species of tick are native to Great Britain (Jameson & Medlock, 2011), 26 to northwestern Europe as a whole (Hillyard, 1996). Most are relatively host specific and primarily nidicolous (i.e. living in or near shelters used by their hosts), and therefore of minimal risk to humans (Gray, Estrada-Pena & Vial, 2014). (Throughout this article we use ‘tick hazard’ to refer to tick species that parasitise humans, and ‘tick risk’ as tick hazard x chance of human exposure).

In contrast to nidicolous species, some ticks feed on diverse host communities, climbing undergrowth or litter and attaching to passing potential vertebrate hosts, including humans. In three regions in England and Wales, patients consulted General Practitioners about tick-bites at a rate of 54–204 per 100 000 inhabitants in 2011, 72.5% of respondents in Cumbria had removed ticks from patients 2011–13 (101/100 000 population) (Gillingham *et al*., 2020). This is only a partial glimpse of the full extent of bites; an estimated ⅓ –⅔ of those fed upon remain unaware (Hofhuis *et al*., 2015), particularly if bitten by smaller instars, and even if noticed many don’t seek medical advice. For example, a 2007 population survey in the Netherlands found a tick bite incidence of 7198/100 000, *c*.1.1 million bites were reported. This equates to approximately fifteen times the number of tick-bite related general practice consultations (Hofhuis *et al*., 2015). Lyme disease is the primary human tick-borne disease of concern in the UK. Cairns *et al*. (2019) used general practice data to estimate a 1-year Lyme disease incidence of 12/100 000 (cautious interpretation is warranted, 59% of these clinical diagnoses lacked documented laboratory confirmation). The causative pathogen of Lyme disease, *Borrelia burgdorferi* s.l., was only identified in 1983 (Sood, O’Connell & Weber, 2011), and over the last decade other human pathogens have been detected in ticks and related hosts in the British Isles, including: spotted fever group rickettsia (Tijsse-Klasen *et al*., 2011; 2013), *Borrelia miyamotoi* (Hansford *et al*., 2015), tick-borne encephalitis virus (Holding, Dowall, Hewson, 2020), and *Babesia venatorum* (Gray *et al*., 2019; Weir *et al*., 2020). In 2019–20 the first presumed autochthonous human cases of tick-borne encephalitis and babesiosis were recorded in the UK (PHE, 2020a). Some of these recently detected health threats may result from emerging foci of imported pathogens. However, it is also possible that in addition to Lyme disease, there may be considerable levels of undiagnosed tick-borne infections affecting persons in the British Isles.

Public Health England have mapped UK tick distributions at 10 km^2^ resolution by combining historical records (Pietzsch *et al*., 2005) with samples sent by the public, who in most cases found them attached to themselves or their pets (Jameson & Medlock, 2011). It should be noted that UK general practice records of arthropod bites do not identify by species (Newitt *et al*., 2016). Hospital Episode Statistics (HES) have been used to map Lyme disease distribution across England (Cooper *et al*., 2017). However, HES uses residential postcodes of patients, not where they were bitten. Thus, whilst HES is valuable to understanding disease burden given that UK tick-borne infections are very often linked to recreational exposure (Dobson, Taylor & Randolph, 2011), the use of this to map differing geographic tick hazard is limited. This is especially true in places such as the SDNP, with high numbers of regional, national, and international visitors. Knowledge on tick density, the most reliable metric of site tick hazard (Ostfeld, 2011), is therefore restricted to the relatively small number of places in Britain actively field-sampled.

The SDNP is a priority area for interventions that reduce tick-borne disease hazard whilst preserving ecosystem health. Prior to our study its downland section had been highlighted by the Health Protection Agency (part-precursor to Public Health England) as a ‘regional foci of Lyme borreliosis’ (HPA, 2012), whilst West Sussex was listed alongside the South Downs as one of 10 areas in England and Wales where Lyme disease infection was most frequent (HPA, 2010). Yet despite this and the Park’s very large visitor numbers, prior to our study no multi-site field sampling of tick hazard in the SDNP, or comparison of hazard between its key habitats, had been published. Elsewhere woodland has been linked to increased tick-borne disease hazard, specifically Lyme disease (Gray *et al*., 1998; Killilea *et al*., 2008), though controversy remains over causal pathways (Levy, 2013). For example, research linking forest fragmentation to increased Lyme disease hazard (summarised best in Ostfeld (2011)) has been criticized by UK researchers (Randolph & Dobson, 2012). Sheep grazing supports vector populations in some UK grass uplands, and though not host competent for *Borrelia burgdorferi* s.l., sheep can support transmission cycles via tick co-feeding (Ogden, Nuttall & Randolph, 1997), and also host *Babesia venatorum* (Gray *et al*., 2019). However, compared to wildlife, the role of livestock in propagation of tick-borne diseases of human concern is under-researched (Stanek *et al*., 2012). Increased wildlife populations have been implicated elsewhere in rising incidence of tick-borne disease (e.g. Crimean-Congo hemorrhagic fever in Turkey (Randolph, 2009a); tick-borne encephalitis in East Europe (Randolph, 2009b)) setting up a potential conflict between biodiversity and human health. Given UK National Parks aim to enhance wildlife and encourage public enjoyment of the countryside (NPUK, 2017), such conflict would be problematic for the South Downs National Park Authority (SDNPA) and the local governments from which most of its members are drawn. However, its joint remit, bioregional framing, and coalition of stakeholder members makes it the ideal body to link and champion site-based and regional policies to reduce hazard, whilst avoiding or reducing conflict between public health and ecosystem health.

### Aims

Our overall project (*Tick-borne hazards in the SDNP and the potential for Planetary Health based interventions*) includes (1) mapping and fieldwork to better understand tick hazard across the SDNP, including crucially at key potential locations for future interventions, (2) a systematic review of proposed interventions to reduce site hazard of the most common tick-borne disease in Britain, Lyme disease, with a focus on those actions not expected to negatively affect ecosystem health. Here we report on our mapping and fieldwork, information on our systematic review can be found in Middleton, Cooper & Rott (2016).

Study objectives:

- identify and describe potential key locations for future interventions;
- map distribution of tick hazard across the SDNP;
- determine tick hazard (species and density) at potential intervention sites; and
- analyse habitat associations with tick hazard in the SDNP.

## MATERIALS AND METHODS

### Sites selection for drag-sampling and potential future interventions

Five sites were selected: three prospectively, and two responsively after submission of ticks obtained by deerstalkers from sentinel deer. The three prospectively chosen sites were located one in each of the SDNP’s three counties. We took this approach so as to sample from along the National Park’s length, and because one of our project’s primary audiences is county authorities which manage countryside sites within the SDNP with high numbers of recreational visitors (e.g. UK accredited country parks as defined by NE & DEFRA (2014)). These authorities are key to implementing potential interventions to reduce tick-borne disease risks in the SDNP as they elect governing members to SDNPA (responsible for strategic action across SDNP), and directly manage downland and woodland sites with high visitor numbers where interventions could be trialed. Of the three counties within SDNP’s borders, two county councils manage such sites in the National Park: Hampshire County Council (SDNP’s western section), and East Sussex County Council (SDNP’s eastern section). West Sussex County Council (SDNP’s central section) does not perform this function within the National Park. The SDNP’s ranger service was consulted about which Hampshire County Council and East Sussex County Council sites had the highest visitor numbers (subsequently confirmed by councils themselves). Given West Sussex County Council do not manage an appropriate site for sampling, a third site was chosen at SDNP’s center which represented a sizeable wealden woodland owned by a key Park stakeholder (Sussex Wildlife Trust).

### Tick collection from Deer

Deerstalkers were recruited through the British Deer Society (bds.org.uk) newsletter and website, sent kits, and asked to collect ticks from deer culled for reasons unrelated to this project. Participants were instructed to inspect the whole animal, collect every visible tick, place them in pre-coded 1ml cryovials (pre-filled with 0.5ml 70% ethanol) and return by post (safety measures, Supplementary Material p. 2). On receipt cryovals were deposited in a laboratory fridge (approx. 5 °C), and after identification transferred to a freezer (approx. -20 °C). Deerstalkers recorded: habitat type; deer species; six-figure grid reference (using ‘OS Locate’ (Ordnance Survey, London, 2014)); body sites ticks found at; and whether ticks were attached or not.

### Tick collection by drag-sampling

Sites were sampled April to November inclusive. To collect questing ticks, sampling was not carried out when air temperature was <7 °C 50 cm above the ground or when vegetation was wet from recent rain/dew, as per James *et al*. (2013). Four sites were sampled in both 2015 and 2016, with an additional site sampled in 2016. At each, six 1m x 50m transects were sampled as per Dobson, Taylor & Randolph (2011). The first two transects chosen were those suspected to have the highest potential exposure of humans to ticks (e.g. vegetation alongside a footpath). Where sites included grassland and woodland, one chosen transect was selected from each. All others were selected using dice and a random number table. To reduce spurious conclusions from single sampling, each transect was planned to be sampled twice yearly, for two years. However, at one private site used for game shooting, it was not possible to visit twice in year-2 due to a requirement to be accompanied by a deerstalker with restricted availability. To improve chances of picking up disease signals in planned follow-up research (usually only a minority of ticks at any site are infected (Vollmer *et al*., 2011)), extras were acquired by drag-sampling between transects, and at follow up visits where possible. Tick sampling techniques differ in efficacy and are affected by habitat/vegetation type (Dantas-Torres *et al*., 2013). To reduce bias ticks were collected simultaneously along transects using woollen blanket, flags, and chaps (*Fig. 1*). Wools were examined after each transect, ticks placed individually in 70% ethanol filled micro-centrifuge tubes, deposited same day in a laboratory freezer (approx. -20°C).

**Figure 1:**
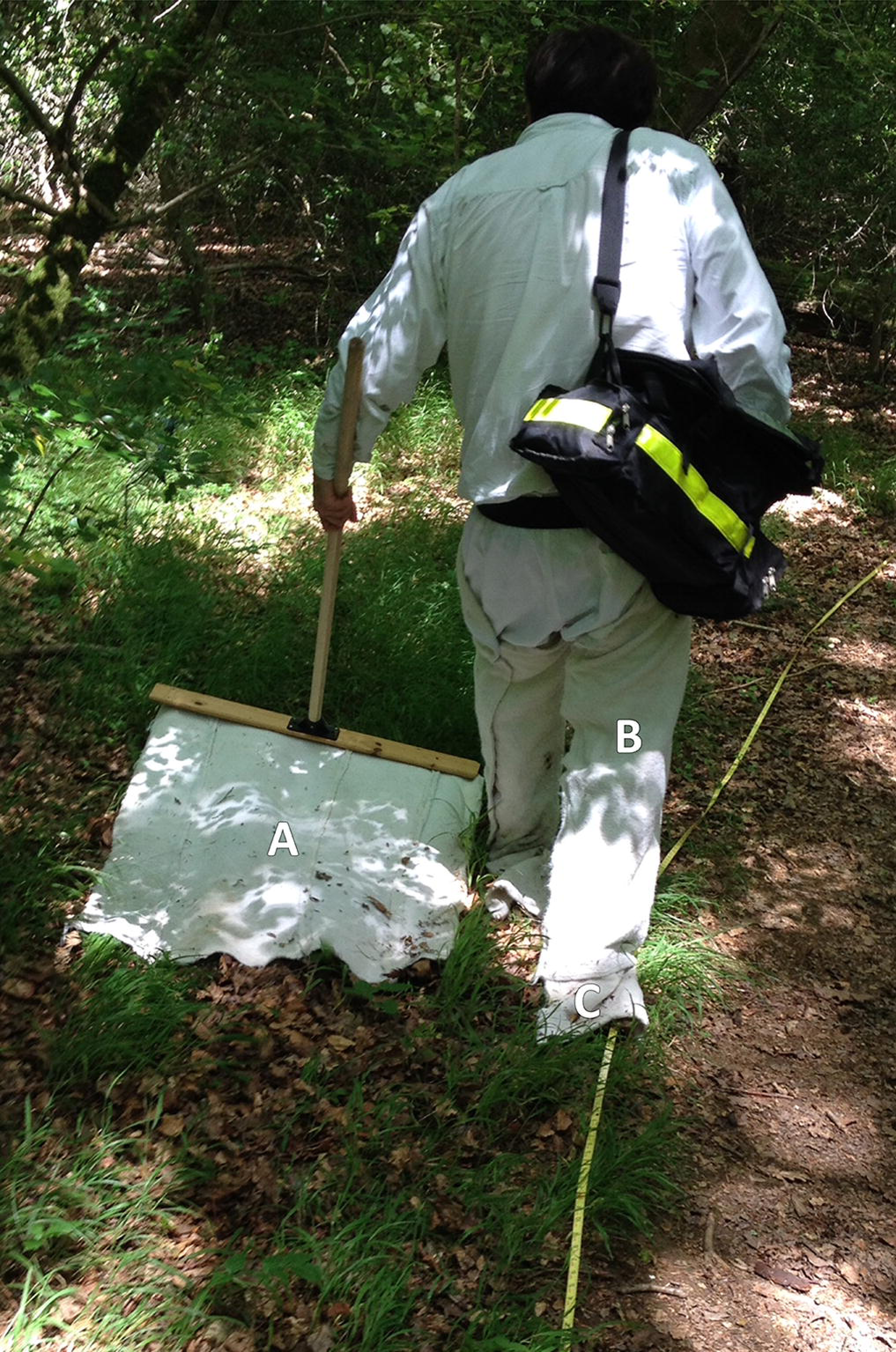
Site tick-sampling equipment. JM drag-sampling along a path border at The Mens, West Sussex. (A) Woollen blanket. (B) Woollen chaps. (C) Woollen flags. Design as per Dobson, Taylor & Randolph (2011). Photo: ASR.

At each transect, at each sampling, photos were taken along with field notes including: ticks collected; date/time; weather; habitat; visitors observed; dominant vegetation; vegetation height; main litter constituents; relative humidity and temperature (both at 50cm and in litter, measured with a Fisher Scientific Traceable Hygrometer). Locations were recorded (10-figure OS grid references, bearings) using ‘OS Locate’ and ‘OS Mapfinder’ (Ordnance Survey, London, 2014) on a Samsung Galaxy Note II phone. (Researcher safety and inter-site contamination control, Supplementary Material p. 2.)

### Tick identification

Identification was conducted in-lab with a hand lens (Hilkinson Ruper x20 15mm achromatic), and where necessary a dissecting microscope (Leica EZ4). A species key was used (Hillyard, 1996), and identification aided by reference to Baker (1999) and Beati, Needham & Klompen (2016). Species and life stage was recorded, adult ticks sexed. Larvae were not fully keyed as clearing for slide mounting would have reduced the sample pool available for future pathogen detection. However, each larva was inspected for characters which identified them to genus, and indicated likely species. If nymphs/adults of more than one species were identified at any site, 10% of larvae from that site (to a maximum of 50) would have been slide mounted and keyed. A limitation of many similar studies has been not enabling retrospective evaluation of species identification (Estrada-Pena *et al*., 2013). To corroborate identification, voucher specimens (including all life stages/sexes) are stored at approx. -20 °C at the University of Brighton, to be deposited into the Natural History Museum acarology collection on publication.

### Mapping distribution of tick hazard

Using ArcMap 10.7 (ESRI, Redlands USA) sites where ticks had been submitted by deerstalkers or drag-sampled by JM were mapped, indicated by points at 100 m^2^ resolution. In addition, to map recorded presence of tick hazard at 10 km^2^ resolution by species, data from the following sources were compared and combined as layers: (1) the most recent published Public Health England/Health Protection Agency tick maps for England and Wales (Cull *et al*., 2018; PHE, 2016; HPA, 2013a, b, c, d), (2) National Biodiversity Network Atlas (NBN, 2020) (which includes historic data 1890 onwards, and into which Public Health England now submits tick records (PHE, 2020b)), (3) a single site drag-sample in 2014 by Layzell *et al*. (2018), (4) point locations of ticks submitted from culled deer or collected by drag-sampling in this project 2015-16, (5) point locations drag-sampled for *Haemaphysalis punctata* by Public Health England and the Animal and Plant Health Agency, primarily 2015-18 (Medlock *et al*., 2018), (6) records from pan-species surveying at Sussex Wildlife Trust reserves, mainly 2016-17 (previously unpublished data). Digital basemaps were obtained from OS OpenData (2020) and Natural England (2020). (Layer generation detailed in Supplementary Material, p. 3.)

### Analysis

As well as site vector presence/absence, for sites drag-sampled both years tick hazard was assessed as (1) questing Density of Ticks, all life stages (DOT), and (2) questing Density of Nymphs (DON). These were calculated as means of totals of four site samplings: six 1m x 50m transects, sampled twice yearly for two years. To determine significance of difference of tick (all life stages) and nymph counts between the four sites sampled in both years, Kruskal-Wallis Tests with follow-on Dunn’s Tests were carried out on counts from all individual transect samplings (i.e. 90 50m x 1m drag-samplings). A further Kruskal-Wallis Test with follow-on Dunn’s Test examined habitat types determining tick hazard. Statistical analysis was carried out in Mintab17 (Minitab Inc, State College, Pennsylvania). Drag-sampling results from the additional site sampled in 2016 are reported as presence/absence, and in distribution mapping, but were not included otherwise in analysis. (Justification of analysis, Supplementary Material p.3.)

### Data availability

All relevant data are included in this article or Supplementary Material (machine readable data deposited at https://sussex.figshare.com/bsms). On publication all novel tick records will be uploaded to NBN Atlas.

## RESULTS

*Figure 2* maps the locations where ticks were submitted from by deerstalkers, and the five potential future intervention sites drag-sampled across the SDNP by the first author. The tick species collected at each site and the locations of nearby towns are also given.

**Figure 2:**
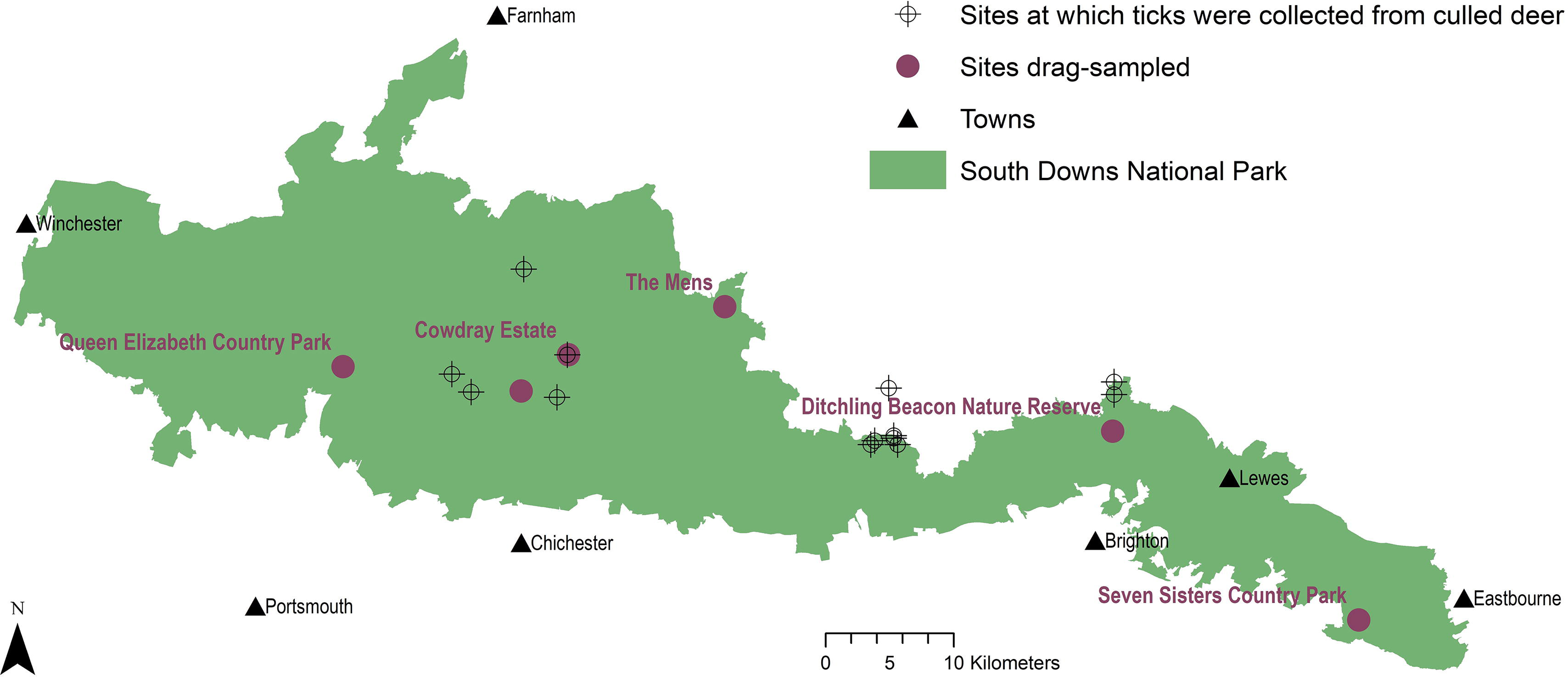
Tick sample collection sites in the South Downs National Park. Sites where ticks had been submitted by deerstalkers or drag-sampled by JM marked by points at 100 m^2^ resolution. All ticks collected from deer were *Ixodes ricinus*, which was also the only tick species drag-sampled at Queen Elizabeth Country Park, Cowdray Estate, and The Mens. *Haemaphysalis punctata* was the only tick drag-sampled at Seven Sisters Country Park and Ditchling Beacon Nature Reserve. Map contains OS data © Crown Copyright (OS OpenData, 2020) and a National Park base layer (unmodified) from Natural England (2020) (https://creativecommons.org/licenses/by-nc-nd/2.0/). Map: JM.

### Sites selected for drag-sampling and potential future interventions

#### Prospectively selected

East Sussex County Council’s site with the highest annual visits (est. 350,000 (ESCC & SDCB, 2004)), was the 280 ha Seven Sisters Country Park (sevensisters.org.uk) in the SDNP’s eastern section. Its visitors centre had 52,124 visitors Jan–Dec 2019 (ESCC, 2020), and only a minority of trips to Seven Sisters Country Park are expected to include a visit to the centre. Unlike the other sites it attracts a large number of international tourists, its white cliffs having featured in major films and as a default Microsoft Windows wallpaper (BBC, 2017; Tsang, 2018; Baddeley, 2020). Seven Sisters Country Park is easily reachable by day-visitors from Eastbourne (8 km away; est. 2019 pop. 114,809 (ONS, 2020)) and Brighton and Hove (24 km away; est. 2019 pop. 244,917 (ONS, 2020)). It is composed of chalk grassland, saltmarsh, shingle seashore, woodland, and a meandering river. Conservation designations include Site of Special Scientific Interest (SSSI), Area of Outstanding Natural Beauty (AONB), Heritage Coast, and Marine Conservation Area. Transects (*Fig. 3A–F*) consisted of sheep grazed chalk downland and woodland, primarily beech (*Fagus sylvatica*) and sycamore (*Acer pseudoplatanus)* (Supplementary Material *Table S1*). Hampshire County Council’s site with the highest annual visits (est. 327,000 (Speller *et al*., 2010)) was the 564 ha Queen Elizabeth Country Park (hants.gov.uk/thingstodo/countryparks/qecp) in the SDNP’s western section. In March 2019– April 20 Queen Elizabeth Country Park’s number plate recognition system recorded 202,559 vehicle entries (HCC, 2020). It is 19 km from Portsmouth (est. 2019 pop. 229,851 (ONS, 2020)), popular with walkers, mountain bikers and picnickers, and hosts outdoor events such as marathons. It consists of downland and wooded hills, designations include: SSSI, National Nature Reserve, Special Area for Conservation, Scheduled Ancient Monuments. All transects (*Fig. 3G–L*) were in woodland (beech, conifer, or hazel (*Corylus avellana*)). Some had sparse undergrowth with dense beech or conifer litter, others nettle patches (*Urtica dioica*) or bramble thickets (*Rubus fructicous* agg.) (Supplementary Material, *Table S2*).

**Figure 3:**
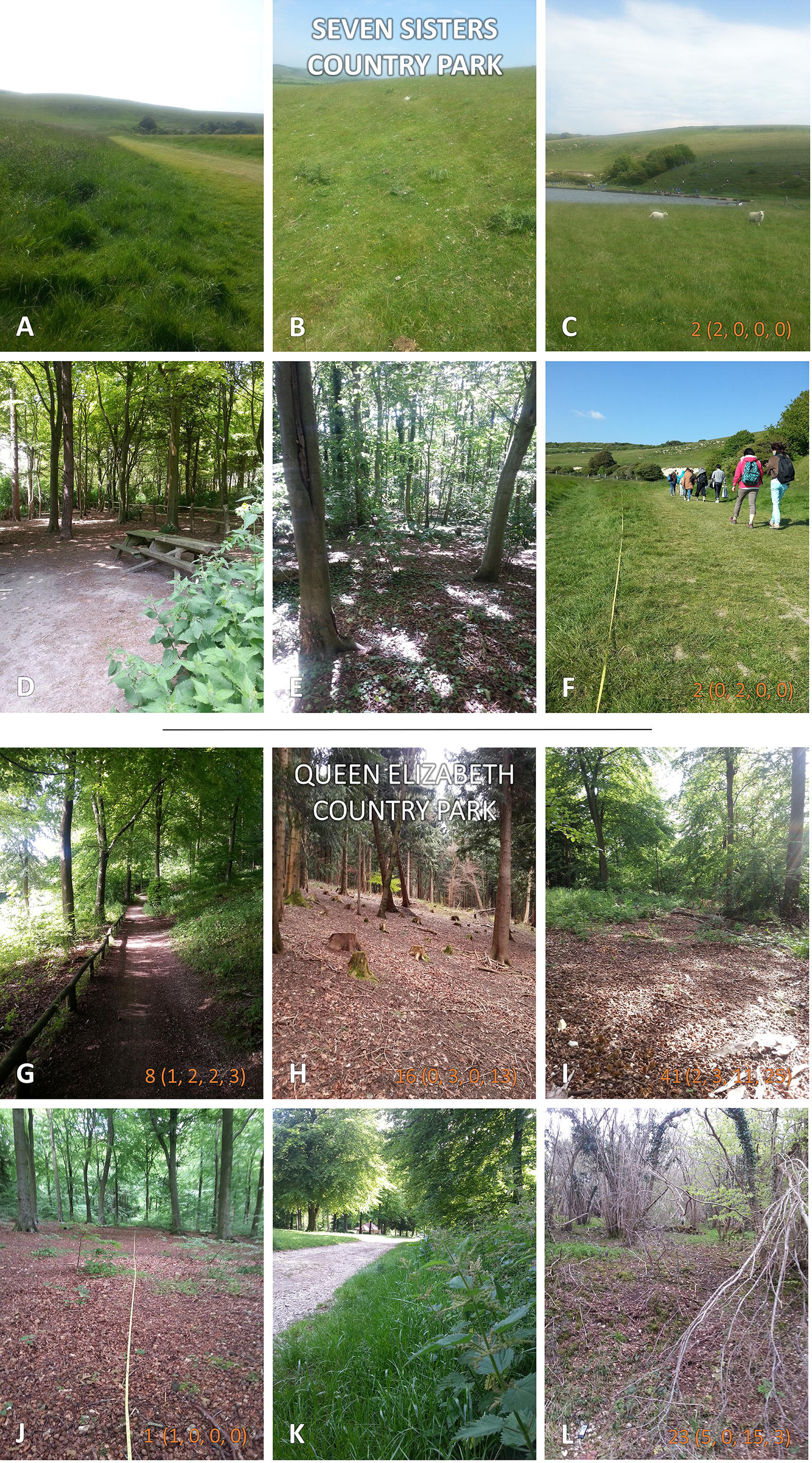
Seven Sisters Country Park, and Queen Elizabeth Country Park. Both sites sampled twice each in 2015 and 2016. Where ticks were present along 50m^2^ transects 2-year totals are given (individual samplings in brackets). Photos: JM.

**Table 1:**
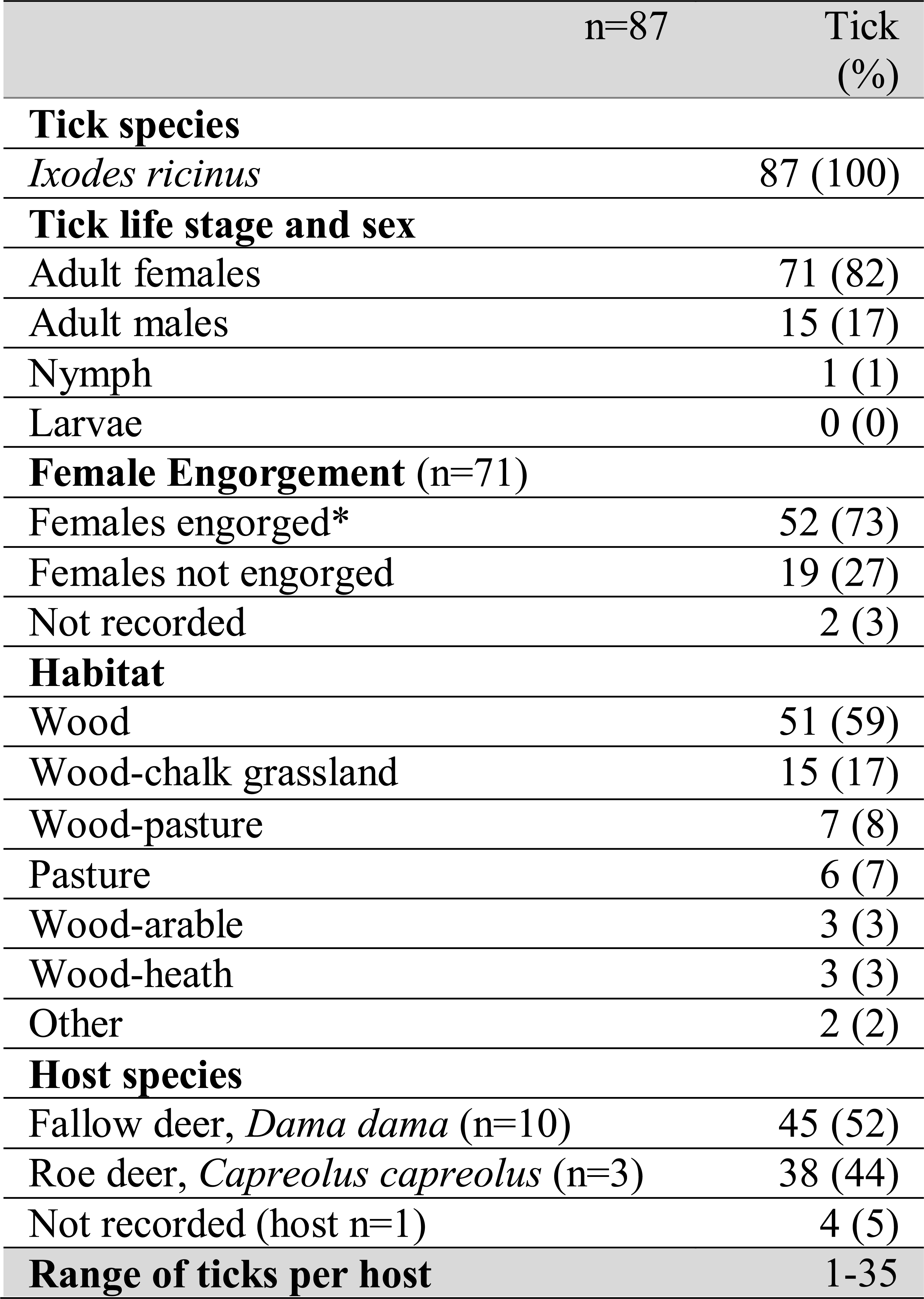
Ticks submitted by deerstalkers. Deer culled for reasons unrelated to this project. Percentages rounded to whole numbers. Engorged status covers adult females and nymphs only, as adult males do not engorge. *in addition the single nymph was engorged.

The third site selected prospectively was woodland at The Mens in West Sussex, a 166 ha nature reserve owned by Sussex Wildlife Trust in the central section of the SDNP (sussexwildlifetrust.org.uk/visit/the-mens). There are less major conurbations close to this section of the SDNP compared to its eastern and western parts. The nearest mid-sized town is Horsham (18 km away; 2011 pop. 49,000 (Horsham District Council, 2016)). The Mens has a small carpark and a network of paths, but is otherwise largely unmanaged wealden forest with relatively few visitors. The site is especially rich in plants, saproxylic invertebrates, and fungi (c.600 species). The sampled transects (*Fig. 4A–F*) followed footpath borders tufted with grass, and cut across ground with sparse undergrowth under high canopies of predominantly beech, and sections with dense waist-high brambles (Supplementary Material, *Table S3*).

**Figure 4:**
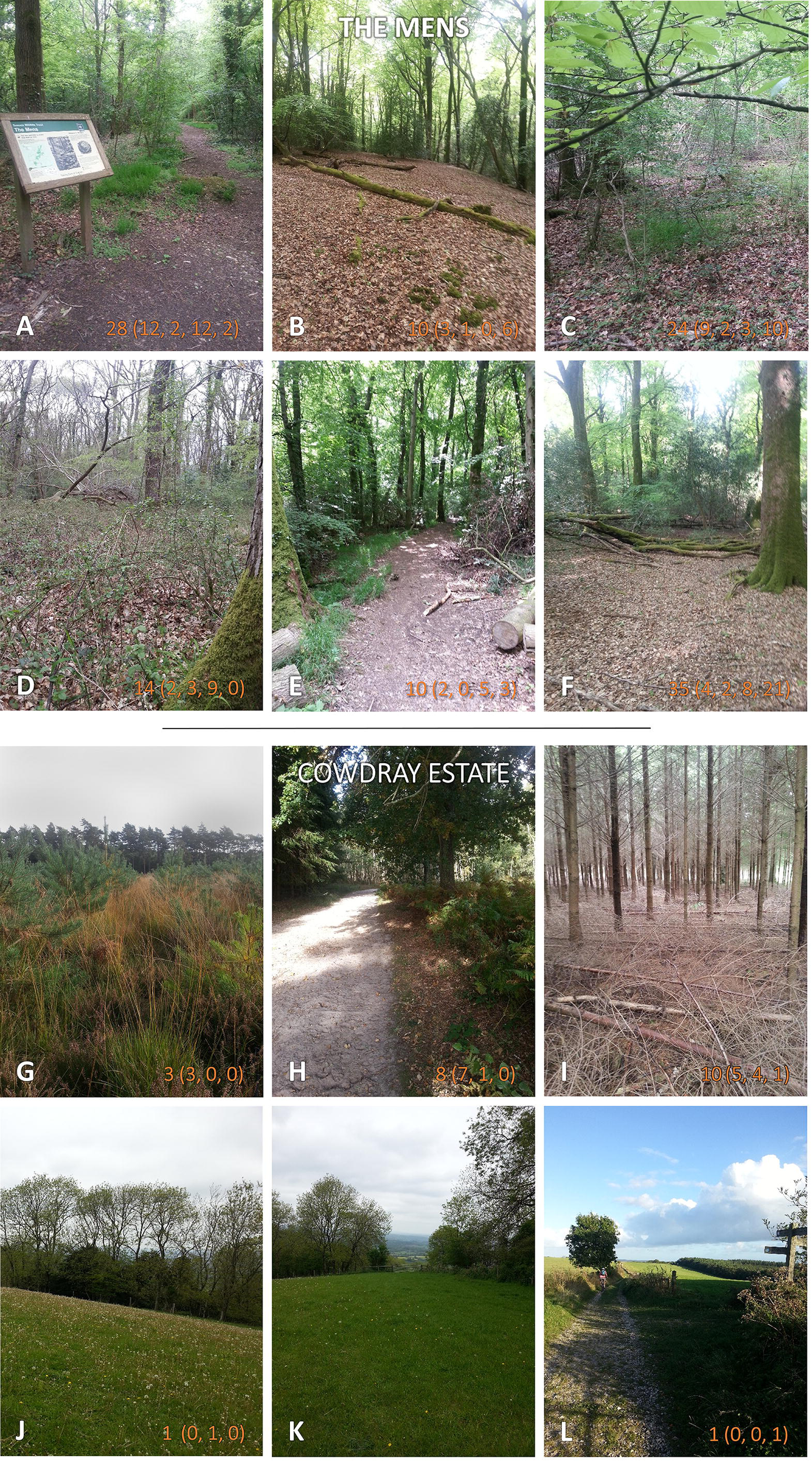
The Mens, and Cowdray Estate. Both sites sampled twice each in 2015 and 2016. Where ticks were present along 50m^2^ transects 2-year totals are given (individual samplings in brackets). Photos: JM.

#### Responsively selected

The 6677 ha Cowdray Estate (cowdray.co.uk) is owned by Viscount Cowdray in the SDNP’s central section in West Sussex, 9 km from Chichester (2011 pop. 26,795 (ONS, 2015)). It is a large private landholding with commercial deerstalking and mostly consists of forestry, downland, arable, and dairy/livestock farming. Cowdray Estate has visitor attractions (golf course, holiday cottages, conference/wedding venue, farm shop and café), and is crossed by well-used public paths. Transects (*Fig. 4G–L*) sampled conifer plantation and sheep-grazed downland (Supplementary Material, *Table S4*). The final site (sampled 2016 only) was the 24 ha Ditching Beacon Nature Reserve in East Sussex (sussexwildlifetrust.org.uk/visit/ditchling-beacon), managed by Sussex Wildlife Trust in the SDNP’s eastern section. Ditching Beacon Nature Reserve is 4 km from Brighton and consists of downland plateau and steep scarp slopes of chalk grassland and woods. The plateau is next to a busy National Trust carpark and is popular with walkers, mountain bikers, and picnickers. Parts of Ditching Beacon Nature Reserve are under conservation grazing with sheep/cattle. The escarpment is an SSSI harboring flower rich chalk grassland, rare orchids, and butterflies. Transects ran through grazed downland, some bordering hawthorn (*Crataegus monogyna*) and ash (*Fraxinus excelsior*) scrub, and along verges of footpaths leading from carparks (Supplementary Material, *Table S5*).

### Extent of tick hazard across the SDNP

#### Distribution and species

Ticks collected by drag-sampling or submitted by deerstalkers confirmed presence across much of SDNP (*Fig. 2*). Ticks were present in both of its characteristic habitats: sheep grazed downland, and the wealden woods. Ticks were found at all sites drag-sampled (though not on all transects, *Figs. 3* and *4*; Supplementary Material, *Tables S1–4*). All ticks submitted from deer were *Ixodes ricinus* (*Fig. 5A*), also the only species collected at three of the four sites drag-sampled in both 2015 and 2016 (Queen Elizabeth Country Park, The Mens, Cowdray Estate). The nationally rare *H.punctata* (*Fig. 5B*) was the sole tick collected from the remaining site sampled in both years (Seven Sisters Country Park), and was also found at Ditchling Beacon Nature Reserve in 2016.

**Figure 5:**
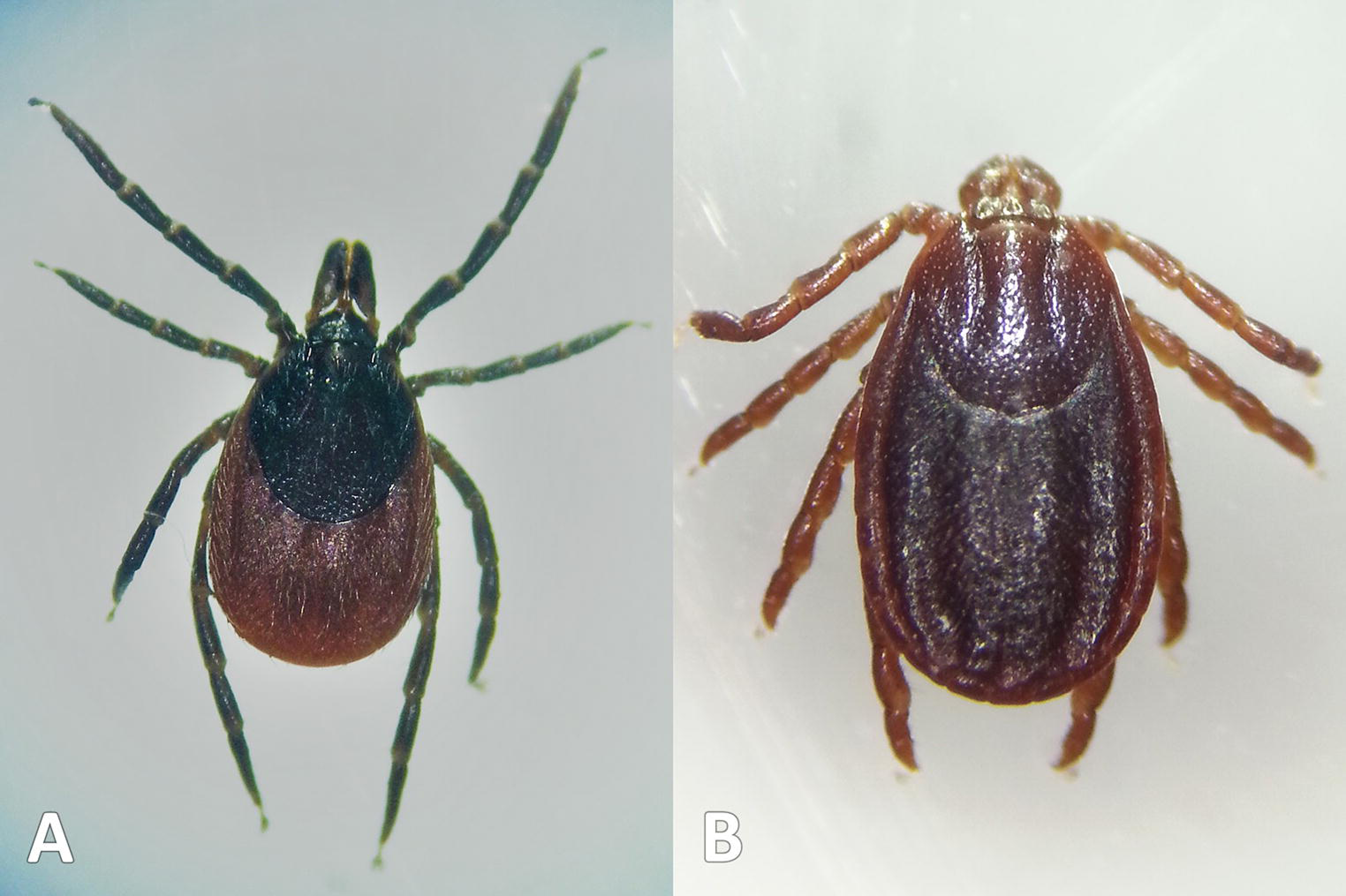
Tick species collected. Ticks collected during study. (A) *Ixodes ricinus* (drag-sampled at The Mens, West Sussex, 2016). (B) *Haemaphysalis punctata* (drag-sampled at Ditchling Beacon Nature Reserve, East Sussex, 2016). Photos: JM.

*Figure 6* maps distribution of tick records at 10 km^2^ resolution. The first report for *I.ricinus*is from 1964, 33/37 of the Parks grid squares have had at least one record (often multiple) in the last 15 years (*Fig. 6A*). In contrast, *Ixodes hexagonus* has been recorded far less, most squares where presence has been recorded represent historic records only (*Fig. 6B*). The earliest *H.punctata* report is from 1920, but all related grid squares have had recorded presence in the last decade, mostly in its known foci in the far east of the Park (*Fig. 6C*). Locations included from recent drag-sampling by Public Health England and Animal and Plant Health Agency suggests it has spread westwards somewhat, and this observation by Medlock *et al*. (2018) is confirmed by drag-sampling in our study at Ditchling Beacon Nature Reserve which extends its known range further still, as does a previously unpublished isolated recording by Sussex Wildlife Trust 44 km further west. A second rare species in the UK, *Dermacentor reticulatus*, was recorded by Sussex Wildlife Trust at one of its West Sussex reserves in 2004 (*Fig. 6D*). To our knowledge *D.reticulatus* has not otherwise been recorded in the SDNP or its constituent counties; this record was not previously included in National Biodiversity Network Atlas or Public Health England/Health Protection Agency published mapping. There are a few records of *Ixodes frontalis* and *Ixodes trianguliceps*. These have not been mapped as they are highly host type specific and not routinely hazardous for humans (Mysterud *et al*., 2015; Drehmann *et al*., 2019).

**Figure 6:**
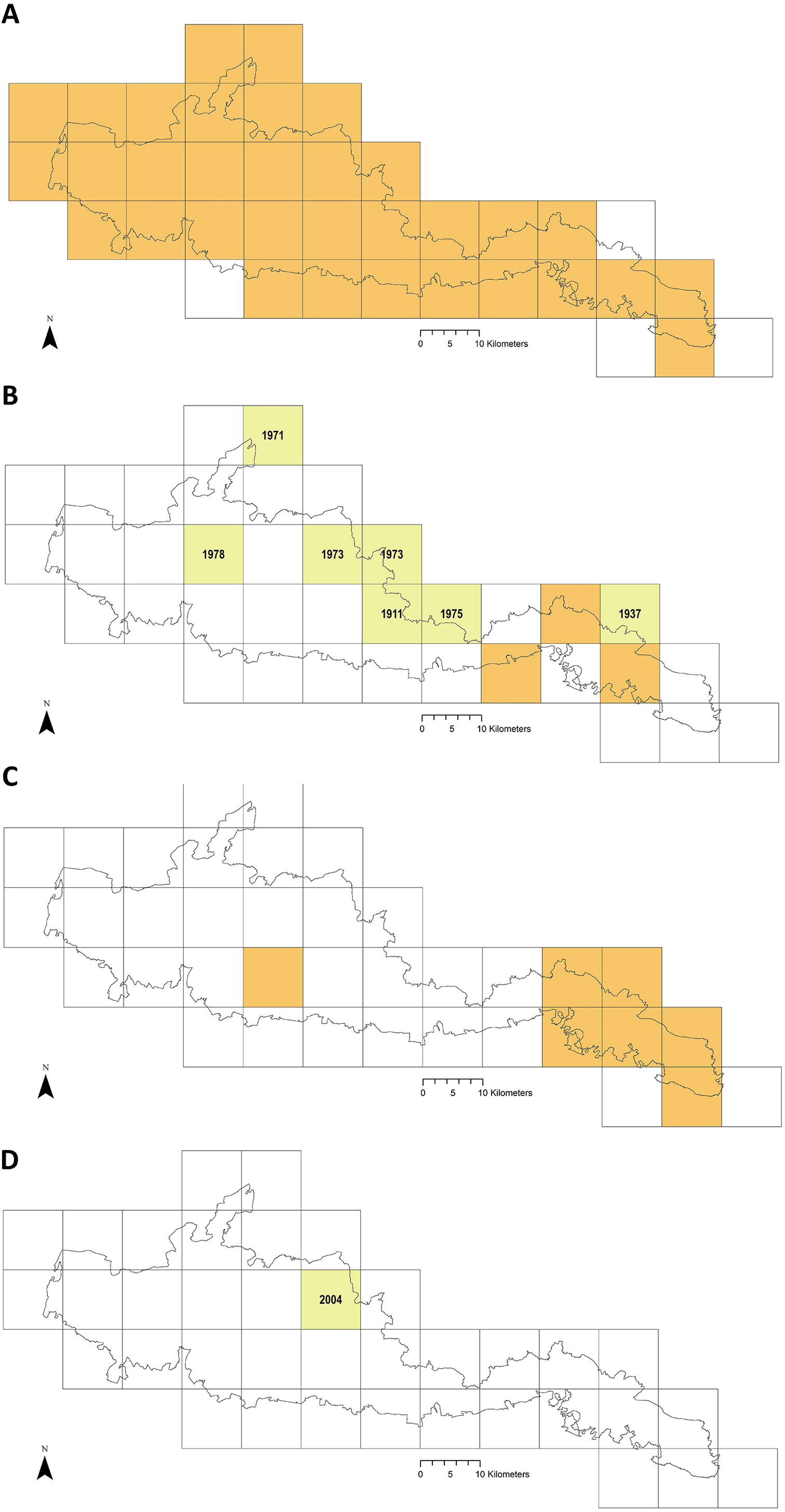
Recorded tick hazard in the South Downs National Park. (A) *Ixodes ricinus*. (B) *Ixodes hexagonus*. (C) *Haemaphysalis punctata*. (D) *Dermacentor reticulatus*. Orange OS grid squares indicate the most recent record/s of tick presence are since 2005 (inclusive). Yellow squares indicate the most recent record/s found were prior to 2005 (latest record date shown). Empty squares represent no records found, but should not be taken as on-the-ground tick absence. Map combines our data of drag-sampling and ticks submitted from culled deer, national maps from the Public Health England/Health Protection Agency tick surveillance scheme (Cull *et al*., 2018; PHE, 2016; HPA, 2013a, 2013b, 2013c, 2013d), the National Biodiversity Network Atlas (NBN, 2020), Medlock *et al*. (2018), Layzell *et al*. (2018), and previously unpublished records from pan-species surveying at Sussex Wildlife Trust reserves. In addition, a case report by Phipps *et al*. (2020) states there were *H.punctata* infestations within the confines of Brighton & Hove in 2019. Maps contain OS data © Crown Copyright (OS OpenData, 2020) and a National Park base layer (unmodified) from Natural England (2020) (https://creativecommons.org/licenses/by-nc-nd/2.0/). Maps: JM.

### Ticks collected from Deer

Eighty-seven ticks were submitted (*Table 1*) obtained from 14 deer at 12 locations (*Fig. 2*). All bar one were adult ticks, and 82% were females. The majority of males were attached in mating; the majority of females were engorged (73%, n=71, two not recorded by collectors), as was the sole nymph. The commonest habitat ticks were collected off deer at was ‘wood’ (59%, n=87). Most of the remaining were from deer shot at mixed edge-habitats involving wood (28%, n=87), e.g. ‘wood-heath’. The majority of hosts were fallow deer (*Dama dama*) (10, 71%), followed by roe (*Capreolus capreolus*) (3, 21%) (n=14, one host species not recorded by collector). Tick burden was 1*–*35 per deer. The most common attachment sites were abdomen and sternum, and posterior and frontal axillae (*Fig. 7*).

**Figure 7:**
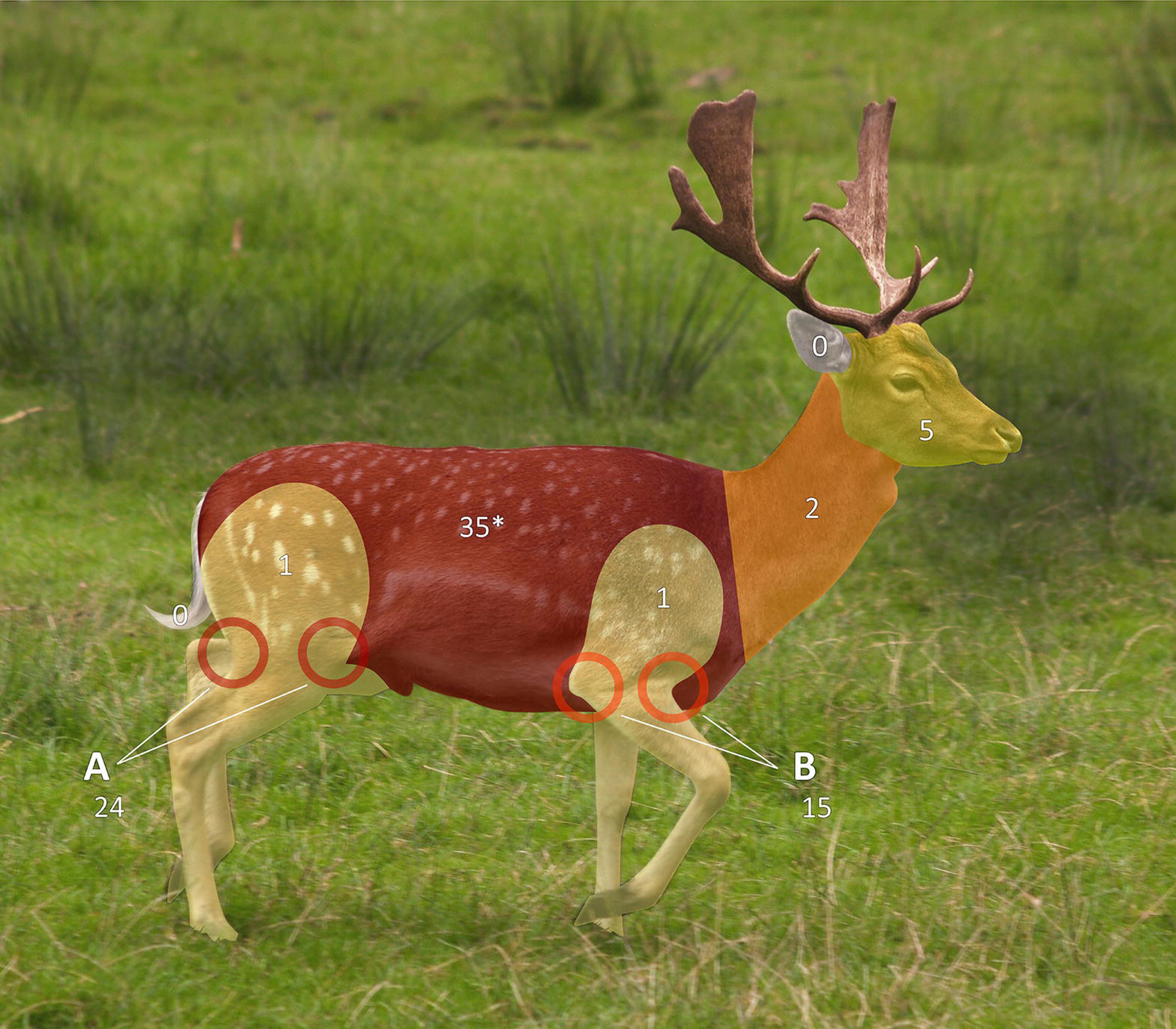
Tick attachment sites on sentinel deer. Ticks collected by British Deer Society members from deer culled for other reasons, attachment sites for four ticks not recorded. Body zones as per Pacilly *et al*. (2014) *Instructions listed abdomen and sternum as separate zones to record, but for 10 ticks this was not done so zones were merged in this figure (reported attachment sites: abdomen, 24; sternum and abdomen, 10; sternum, 1). Photo: Johann-Nikolaus Andreae (2008), use and changes made under CC-BY-SA-2.0 which also applies to this figure. Original: https://web.archive.org/web/20200930080151/ https://commons.wikimedia.org/wiki/File:Fallow_deer_in_field_%28cropped%29.jpg.

### Ticks collected by drag-sampling

Drag-sampling four sites in both 2015 and 2016 collected 622 ticks (*Table 2*). Of these, 237 were along transects and are included in calculations of vector densities and analysed statistically. 385 extras were stockpiled to aid future pathogen detection. Ticks were present at all four sites and the additional site sampled twice in 2016 (1 adult only). Of ticks collected by drag-sampling at all sites (n=623), most were nymphs (53.8%), followed by larvae (42.5%), and a small number of adults (3.7%, 13 males, 10 females (Supplementary Material, *Tables S1–4*). 93.3% (222) of ticks gathered along transects (n=238) had attached to woollen blankets, 6.7% (16) to woollen chaps, and none were attached to flags (*Table 2*). *Fig*ure 8 shows a breakdown of tick data from transects by collection month, and by tick life stage (all sites for y1 and y2, n=238).

**Figure 8:**
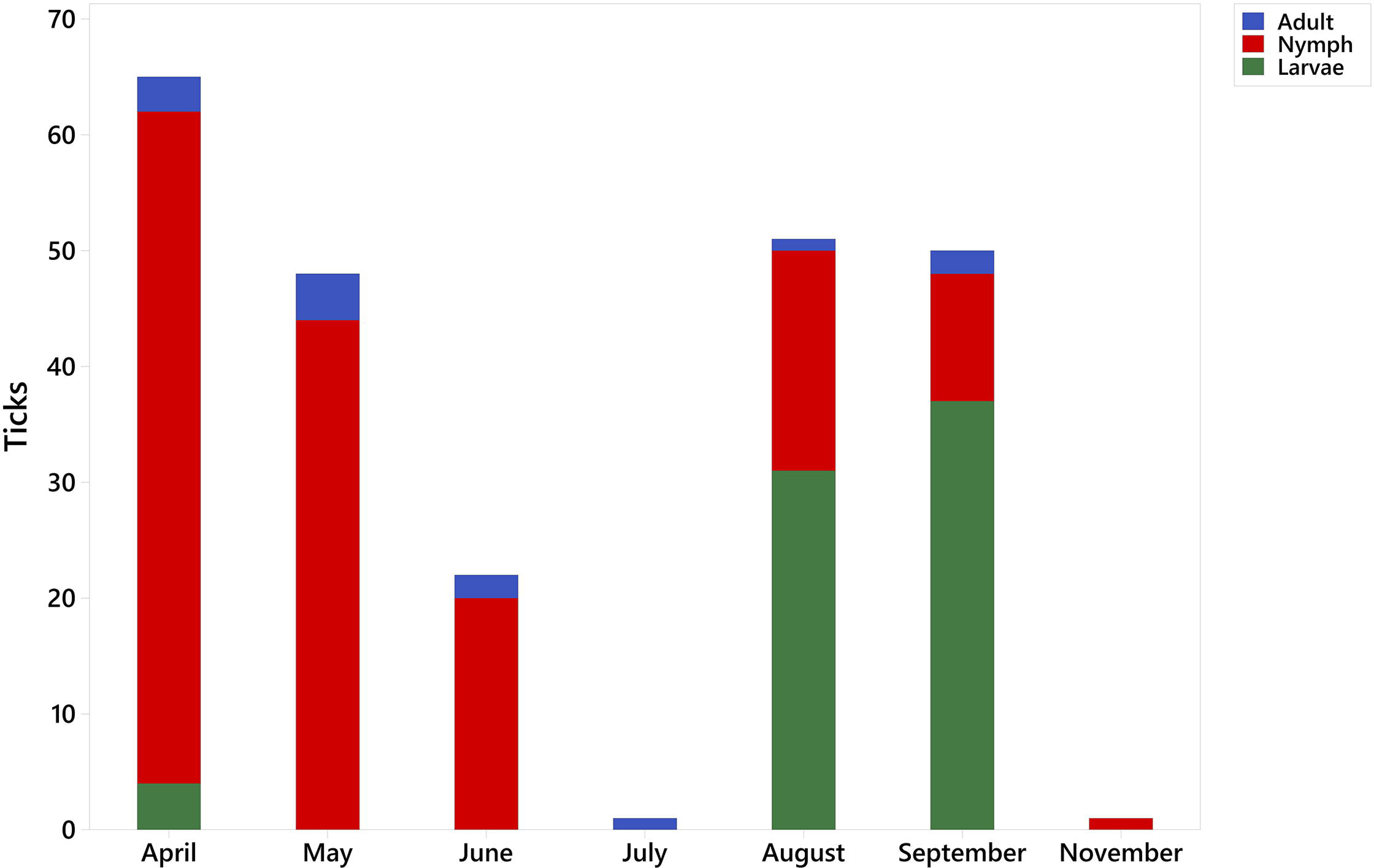
Life stages of ticks collected along study transects by month. 2015 and 2016: Seven Sisters Country Park, Queen Elizabeth Country Park, The Mens, Cowdray Estate. 2016 only: Ditchling Beacon Nature Reserve. Nymphal and larval proportions in sampling months partly reflect *Ixodes ricinus* annual lifecycles, and are not presented to indicate changing *quantity* of hazard, as monthly total differences may be partly explained by which sites, and how many, were visited.

**Table 2:**
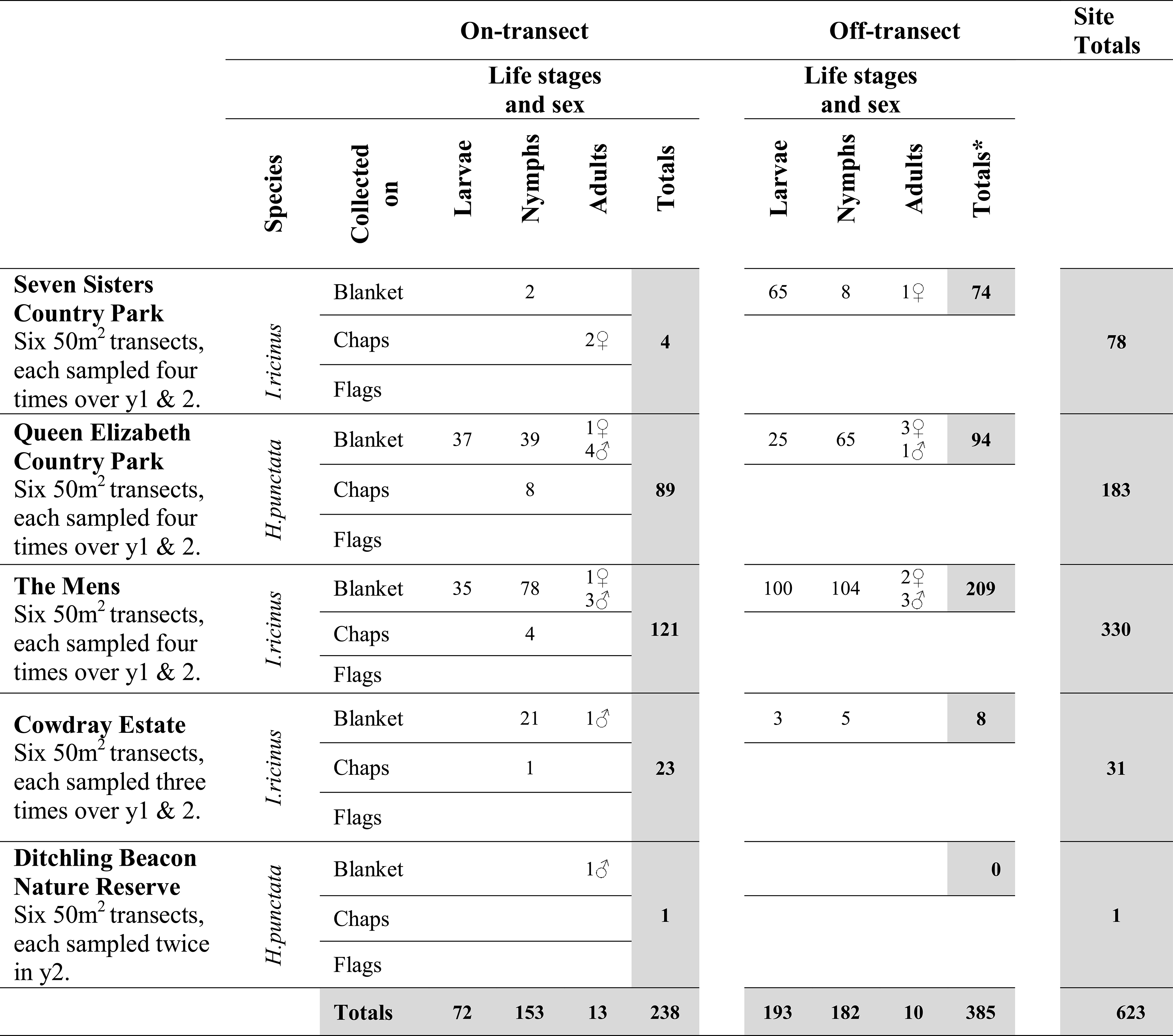
Ticks collected by site drag sampling. Results of individual samplings in Supplementary Material, Tables S1-4.

**Table 3:** Tick hazard and visitors observed during transect drag sampling. Results of individual samplings in Supplementary Material, Tables S1-4.

|  | Visitor numbers observed | Tick species | Ticks found on transects | Ticks per individual 50 m<sup>2</sup> sampling | Site Density of Nymphs (DON) | Site Density of Ticks (DOT) |
| --- | --- | --- | --- | --- | --- | --- |
| The Mens | Low to none | *I. ricinus* | 121 | 0–21 | 30/300 m<sup>2</sup> | 30/300 m<sup>2</sup> |
| Queen Elizabeth Country Park | Very high | *I. ricinus* | 89 | 0–25 | 12/300 m<sup>2</sup> | 22/300 m<sup>2</sup> |
| Cowdray Estate | Low | *I. ricinus* | 23 | 0–7 | 6/300 m<sup>2</sup> | 8/300 m<sup>2</sup> |
| Seven Sisters Country Park | Very high | *H. punctata* | 4 | 0–2 | 1/300 m<sup>2</sup> | 1/300 m<sup>2</sup> |
| Ditchling Beacon | Moderate | *H. punctata* | 1 | 0–1 | n/a | n/a |

### Tick hazard at potential sites for interventions

#### Site-by-site

Seven Sisters Country Park: *Figures 3A–F* show the number of ticks obtained at each transect, at each sampling (Supplementary Material, *Table S1*). 78 *H.punctata* ticks were collected, four along transects, 74 off-transect, representing all life stages, attached to both blanket (76) and chaps (2) (*Table 2*). On-transect ticks were nymphs (2) and adults (2). All ticks were collected on downland, none having been collected on the wooded transects (Supplementary Material*, Table S1*). Tick hazard ranged 0–2 ticks per 50 m^2^ individual sampling, site DON and DOT were both 1 per 300 m^2^ (*Table 3)*. There were hundreds of visitors observed every day. The area which provided most off-transect ticks is in the background of the photo of transect 3 (*Fig. 3C*), hosting a school picnic.

Queen Elizabeth Country Park: *Figures* 3G–L show the number of ticks obtained at each transect, at each sampling (Supplementary Material, *Table S2*). 183 *I.ricinus* ticks were collected, 89 on-transect, 94 off-transect, from all life stages, attached to both blanket (175) and chaps (8) (*Table 2*). On-transect ticks were larvae (37), nymphs (47), and adults (5). Ticks were found on five of the six transects, all of which were wooded or on paths verging woodland (Supplementary Material*, Table S2*), range 0–25 per 50 m^2^ sampling, DON=12 per 300 m^2^, DOT=22 per 300 m^2^ (*Table 3*). A very large number of visitors were present during all site visits, and though some transects were especially busy (e.g. *Fig. 2G*, running from the visitor centre) people were observed during sampling at all transects, even far from carparks.

The Mens: *Figures* 4A–F show the number of ticks obtained at each transect, at each sampling (Supplementary Material, *Table S3*). 330 *I.ricinus* ticks were collected, 121 on-transect, 209 off-transect, representing all life stages, attached to both blanket (296) and chaps (4) (*Table 2*). On-transect ticks were larvae (35), nymphs (82) and adults (4). Ticks were obtained from all six transects (Supplementary Material, *Table S3*), range 0–21 per 50 m^2^ sampling, DON=21 per 300 m^2^, DOT=30 per 300 m^2^ (*Table 3*). Few visitors were met during sampling visits and, unlike any of the other sites, sometimes none were encountered.

Cowdray Estate: *Figures* 4G–L show the number of ticks obtained at each transect, at each sampling (Supplementary Material, *Table S4)*. 31 *I.ricinus* ticks were collected, 23 along transects, eight off-transect, from all life stages, attached to both blanket (29) and chaps (1) (*Table 2*). On-transect ticks were nymphs (22) and adult (1). Ticks were collected on five of six transects, on both woodland and downland (Supplementary Material*, Table S4*), range 0–7 per 50 m^2^ sampling, DON=6 per 300 m^2^, DOT=8 per 300 m^2^ (*Table 3*). Public footpaths along transects (*Fig.4H* and *L*) were in use by walkers during all visits. One of two downland transects on which a tick was found ran along the South Downs Way.

Ditchling Beacon Nature Reserve: One single *H.punctata* tick was collected on-transect on semi-grazed downland adjacent to scrub woodland, none on the other five transects at either set of samplings in 2016. The adult male tick had attached to the blanket (*Table 2*). Densities were not calculated due to 1-year only sampling (Supplementary Material, p. 3). Every day during sampling large numbers of visitors were observed crossing the high plateau, few on the lower scarp slope where the tick was collected.

### Comparisons between sites

Tick hazard was detected at all four sites surveyed in both 2015 and 2016, but levels (range per sampling, DON, DOT) differed between them, as outlined above and shown in *Table 3.* Kruskal-Wallis Tests showed average ranks for (i) nymphs, and (ii) ticks (all life stages) differed significantly (<0.05) for at least one of the four sites (nymphs: H=31.59 (DF 3, n=90), p=0.000 (adjusted for ties); ticks (all life stages): H=31.93 (DF 3, n=90), p=0.000 (adjusted for ties)). Post-hoc Dunn’s Tests were used to carry out pairwise site comparisons (magnitudes and directions of differences, Supplementary Material *Fig. S1*). The Mens had significantly more nymphs than Cowdray Estate (Z=2.128, p=0.0000), Seven Sisters Country Park (p=0.0005), and Queen Elizabeth Country Park (p=0.0121). Seven Sisters Country Park had significantly less than Queen Elizabeth Country Park (p=0.0029). For ticks (all life stages) The Mens had significantly more ticks than Cowdray Estate (Z=2.128, p=0.0013) and Seven Sisters Country Park (p=0.000) which in turn had significantly less ticks than Queen Elizabeth Country Park (p=0.006). However, unlike for nymphs, The Mens did not have significantly more ticks (all life stages) than Queen Elizabeth Country Park.

### Habitat associations with tick hazard in the SDNP

Of the four sites drag-sampled in both years, sites with transects entirely in woodland (Queen Elizabeth Country Park, The Mens) had the highest tick hazards (*Tables 2 and 3;* Supplementary Material*, Tables S2* and *S3*). However, tick hazard was present at all sites, including on the grazed downland sections of the two sites (Seven Sisters Country Park and Cowdray Estate) which had transects in downland and woodland (Supplementary Material*, Tables S1* and *S4*). The tick hazard present was not universal in wooded sections of sites; no ticks were found in the forested part of Seven Sisters Country Park (Supplementary Material*, Table S1*). A Kruskal-Wallis Test performed on multiyear means of ticks collected (all life stages) on each habitat coded transect showed average ranks differed significantly (<0.05) for at least one of the three coded habitat types (H=6.39 (DF 2, n=24), p=0.041 (adjusted for ties)). A post-hoc Dunn’s Test was used to carry out pairwise habitat type comparisons (magnitudes and directions of differences, Supplementary Material *Fig. S2*). There was not a statistically significant difference in the number of ticks (all life stages) between deciduous woodland and conifer woodland/planting. However, both these habitats had significantly more ticks than downland (Z=1.834, p=0.0226, and p=0.0390 respectively).

## DISCUSSION

*Ixodes ricinus* or *H.punctata* ticks were recorded at all sites drag-sampled, on some transects in high numbers (*Fig. 3* and *4)*. The extent of tick hazard differed between sites, and was significantly higher in woodland compared to grazed downland. Although it must be noted that tick hazard was still present at all downland sites. The mapping presented in this paper indicates that tick hazard is widely distributed across the SDNP, as confirmed by the deerstalker submission of ticks from sites across the National Park.

*Ixodes ricinus* (*Fig. 5A*) is the tick most often affecting humans and pets in the UK (Jameson & Medlock, 2011; Abdullah *et al*., 2016; Davies *et al*., 2017). It is therefore unsurprising it was the species most frequently recovered in the drag-sampling and deerstalker submissions, and the most spatially reported (*Fig. 6*). The larvae and nymphs of this species feed primarily on rodents and small birds, while the adults mainly parasitise larger mammals. Transovarial transmission of some pathogens can sometimes cause larvae to hatch as infectious (Hauck *et al*., 2020). However, a nymph feeding on a human will also have had opportunity to become infected when feeding as larva, and an adult will have had two blood meals, potentially from very different animals. Thus *I.ricinus* can act as a vector for the transmission of pathogens to humans from diverse taxa (Hillyard, 1996; Randolph, 2009b; Mannelli *et al*., 2012). It is the UK’s most common Lyme disease vector, followed by *I.hexagonus* (Jameson & Medlock, 2011; Medlock & Leach, 2015), and based upon the data presented here from the drag-sampling and GIS mapping, it is likely to be responsible for the majority of Lyme disease cases contracted in the SDNP. *Ixodes ricinus* has also been reported to be Europe’s major tick borne encephalitis vector (Brugger *et al*., 2017). Tick borne encephalitis virus has been detected in one of the Park’s host counties, but to our knowledge no ticks within the SDNP have been tested. Layzell *et al*. (2018) drag-sampled a single site within the National Park in 2014 (West Dene, West Sussex) and isolated *Borellia miyamotoi* in *I.ricinus* ticks from that site. Like Lyme disease, *Borellia miyamotoi*disease is caused by *Borrelia* species, but signs and symptoms markedly differ so that it is classed separately (Telford *et al*., 2015). In the USA, a case series of 94 individuals (identified by retrospectively testing stored patient samples) indicated a clinical presentation of chills, headache, generalised/joint pain, thrombocytopenia, and high fever. Of these people, 24% of cases required hospitalisation, and all responded well to antibiotics (Molloy *et al*., 2015). *Borellia miyamotoi* disease was discovered far more recently than Lyme disease (the first confirmed Western European case was in 2013 (Fonville *et al*., 2014)). A clear picture of disease burden in humans is, therefore, not yet available. *Borellia miyamotoi* detection in the SDNP, in what we found to be the Park’s most well distributed tick vector, adds further weight to the need to conduct interventions.

UK countryside workers perceive spatial overlaps between widening deer abundance and *I.ricinus* (Scharlemann *et al*., 2008), but wider ecological determinants such as host community compositions affect densities of infected ticks and thus actual disease hazard and determinants vary between site (Kurtenbach *et al*., 1998; Keesing *et al*., 2010). For instance, whilst deer likely have roles in most, but not all, UK Lyme disease systems (Ogden *et al*., 1997; Gilbert *et al*., 2012) they are non-competent hosts for the pathogen; small mammals/birds are usually required as disease reservoirs (Franke, Hilebrandt & Dorn, 2013). *Ixodes ricinus* is often associated with forests (Ehrmann *et al*, 2017), and in our fieldwork its presence and densities were highest in wooded areas. However, we also collected the species on sheep-grazed land (as elsewhere in the UK (Evans, Sheals & Macfarlane, 1968; Ogden *et al*., 1997; Gilbert *et al*., 2017)). Management strategies across the SDNP should take this into account, especially where downland is bounded by woods (Gilbert *et al*., 2017).

*Haemaphysalis punctata’s* (*Fig. 5B*) continued and expanding presence in the SDNP is evident in our tick hazard mapping (*Fig. 6C*), raising concerns about pathogens it can vector, including *B.burgdorferi* s.l. (Tälleklint, 1996), tick-borne encephalitis virus (Estrada-Peña & Jongejan, 1999), and spotted fever group rickettsiae. UK *H.punctata* testing has so far been negative for *B.burgdorferi* s.l. (Tijsse-Klasen *et al*., 2013), to our knowledge un-conducted for tick-borne encephalitis virus (known ranges do not presently overlap), but positive for spotted fever group rickettsiae at some sites outside the Park (Tijsse-Klasen *et al*., 2013). Spotted fever group rickettsiae are an emerging European disease threat (Lindblom *et al*., 2013), but one that may have been present yet unidentified for some time (Vitale *et al*., 2006). For example, *Rickettsia massiliae* was first identified as a human pathogen in 2005 after isolation from a clinical sample collected 20 years prior (Vitale *et al*., 2006). Spotted fever group rickettsiae related misdiagnoses and under-reporting still likely continue (Tijsse-Klasen *et al*., 2013). *Haemaphysalis punctata* is known to parasitise humans (Hillyard, 1996) and tick submissions to Public Health England by the public show this is happening in the SDNP (Medlock *et al*., 2018; Phipps, 2019). Sheep and cattle are its main adult hosts, others include horses, hedgehogs, rabbits, birds, goats, deer, and mustelids (Evans, Sheals & Macfarlane, 1968; Hillyard, 1996). Despite flock treatments, sheep infestation at Seven Sisters Country Park (one of the sites we collected it at) has been present for decades (personal communication with site sheep farmer, 2015; Medlock *et al*., 2018). In 2020, 11.5% of a sheep flock in the SDNP near Lewes suffered fatal tick pyaemia, the first such UK outbreak connected to *H.punctata* (Macrelli *et al*., 2020). Also in 2020, on Brighton’s downland outskirts (near the second site we collected *H.punctata*) sheep used for conservation grazing had to be removed on welfare grounds following heavy infestations with the tick (Phipps *et al*., 2020). Given these and related incidents Animal Plant Health Agency and Public Health England are investigating further in the Park and working with farmers.

Medlock *et al*. (2018) state that there does not generally seem to be habitat overlap in the UK between *H.punctata* and *I.ricinus*. Though *Fig.6A* and *C* show that in all 10 km^2^ OS grid squares where *H.punctata* have been detected *I.ricinus* has also been recorded, this is not in fact in contradiction. On a finer scale of the individual sites JM drag-sampled, tick presence was either *H.punctata* or *I.ricinus*, not both. *I.ricinus* grassland preference in the UK for rough grazing is likely connected to the rapid desiccation it experiences in short grass lacking thick mats of vegetation and litter (Evans, Sheals & Macfarlane, 1968). In contrast, *H.punctata* can likely survive better in short grass, its traditional range includes deserts (Nosek, 1973). Outside the UK *H.punctata* is also found in forest (Bor an *et al*., 2020). Whilst its established foci in the eastern Downs is relatively unwooded, if allowed to expand its range westward along the downland ridge it will increasingly encounter patchworks of grazing, scrub, and woods. The two species site occupancy and host community may then begin to overlap, with implications for pathogen carry and thus hazard to humans.

### Recommendations for key locations for future interventions

Queen Elizabeth Country Park: Given its high tick hazard and very high annual visitor numbers this site is the highest priority for interventions. It also lays in Hampshire where tick-borne encephalitis virus has been detected and would be a logistically simple trial setting for action which may be required elsewhere in the county. Basic measures, such as increased frequency of mowing verges (Medlock *et al*., 2012; Del Fabbro, 2015) and leaf litter removal (Schulze, Jordan & Hung, 1995) would be implausible across the whole Country Park, but could reduce contact at small high risk plots: e.g. edges of marked picnic areas; such as along the path that goes past the visitor centre (*Fig. 3G)*. It hosts large outdoor sports events, and elsewhere tick removal/submission from participants has been used to tick-sample (Hall *et al*., 2017). Site sampling by this method would be inexpensive, and along with increased signage would raise awareness of tick presence and the value of carrying out post-activity tick-checks. These are important as early tick removal reduces transmission, and during-activity recommendations aimed at individuals to minimise exposure are unlikely to be headed (Middleton, Cooper & Rott, 2016).

The Mens: The site had the greatest DOT and DON, but fewest visitors. Thus though its DOT was thirty times Seven Sisters Country Park’s (annual visitors, est. >300,000 (ESCC and SDCB, 2004)), the tick risk to public health is far smaller. Nevertheless, signage in the carpark would be beneficial and the site could become a useful research/trial location. It’s impressive beach masts (the dominant litter constituent of most of its transects) may support high rodent densities, which can be very host-competent for ticks and pathogens, amplifying disease hazard (Keesing *et al*., 2009; Ostfeld *et al*., 2014; Krawcyk et al., 2020). Such a relationship has been observed in northeastern USA where high acorn masts cause subsequent year surges in rodents, followed by elevated Densities of Infected Nymphs (DIN) (Ostfeld *et al*., 2006). Predator protection and reintroduction have been proposed as ecologically beneficial interventions (Nilsen *et al*. 2007; Levi *et al*., 2012). For example, Hofmeester *et al*. (2017) observed an indirect negative correlation of red fox and stone marten activity with DON and DIN at forest plots across the Netherlands, and called for wider predator appreciation and protection. Tick pathogen testing could establish if the very high tick densities at The Mens are matched by high densities of infection. If so, measurement of vertebrate, especially rodent, tick and pathogen burden would determine if predator re-introduction (i.e. pine martens) and protection (i.e. foxes) should be trialed as health interventions at the site and environs. If successful, this could be extended further through the wooded wealden section of the SDNP of which it is a part. Given The Mens is a relatively large biodiverse reserve for South-East England (Whitbread, 2013), if infection densities are lower than expected based on tick densities, explanatory work could contribute to scientific debate on relationships between biodiversity and health (see: Randolph & Dobson, 2012; Levy, 2013; Foley & Piovia-Scott, 2014; Civitello *et al*., 2015).

Cowdray Estate and Deerstalkers: With comparably low tick hazard and fewer visitors than most other sites, the Estate itself does not need to carry out interventions to reduce site tick hazard. However, action aimed at walkers crossing Cowdray Estate and deerstalkers working in it (and by implication elsewhere under similar circumstances) is warranted. Publicly/NGO maintained car parks along the route of the South Downs Way in Cowdray Estate would benefit from tick related signage (absent during our visits). To our knowledge no study of *B.burgdorferi* s.l. seropositivity or Lyme disease cases has been conducted for professional UK deerstalkers as has been for more commonly considered occupationally at risk groups such as foresters (e.g. De Keukeleire *et al*., 2018). However, British Deer Society volunteers at multiple sites stated to the lead author of this paper that during stalking and butchery they regularly encounter ticks, often getting bitten. This statement is plausible given data presented in *Table* 1 and *Fig.* 7. On a precautionary principle, professional deerstalkers would thus benefit from provision of permethrin-impregnated clothing which is effective against ticks (Faulde *et al*., 2015), and targeted tick-related education. They would also be a priority group for future tick-related disease vaccination efforts. A vaccine against European *B.burgdorferi* s.l. strains is under development (Nayak *et al*., 2020), and one for tick-borne encephalitis is available but only recommended at present for those doing outdoor activities in a country where tick-borne encephalitis virus is common (NHS, 2021).

Seven Sisters Country Park and Ditchling Beacon Nature Reserve*: Haemaphysalis punctata’s* original distribution at Cuckmere Haven and surroundings suggests importation via migratory birds (Tijsse-Klasen *et al*., 2013), but how it is spreading westward is unclear. This may be facilitated by livestock movements (including conservation grazing (Medlock *et al*., 2018)), birds (immature tick stages), or pets (Public Health England has received submissions taken off dogs locally (Phipps, 2019)). Public Health England and Animal Plant Health Authority are carrying out targeted surveys to determine its invasion boundary (*Fig. 6C*) and means of spread, knowledge required for effective region-level intervention. Additional work to understand its ecology and control is needed (Medlock *et al*., 2018). Acaricide application to sheep on the Lewes site resolved the situation in-year (Phipps *et al.,* 2020). However, as the data from Seven Sisters Country Park indicates, livestock treatment alone may be insufficient, potentially because *H*.*punctata* instars are supported by non-domestic hosts. Pasture spelling (i.e. temporary flock removal for c.6 months) is a traditional practice to reduce tick numbers (Hillyard, 1996), yet may be unsuccessful in clearing *H*.*punctata* from grazing land given unfed individuals can survive relatively long periods without blood meals: nymphs, 252 days; adults, 255 (Evans, Sheals & Macfarlane, 1968). Trials at Seven Sisters Country Park, Ditchling Beacon Nature Reserve, and related sites could evaluate approaches for control whilst reducing on-site tick hazards. No tick-related notices were seen during sampling or subsequent visits (up to January 2021) at either site. Instructive, but not alarming, signs could be placed in carparks at both, as has been done unobtrusively at another site, Mount Caburn, which is a less visited local site, but where *H*.*punctata* is also present. Notices should emphasize strongly the need for post visit checks, including vigilance over longer subsequent periods than normally recommended (*H.punctata* nymphs usually feed for one week, but may attach up to 33 days (Evans, Sheals & Macfarlane, 1968)).

### Strengths and weaknesses

This work is part of a regional intervention planning exercise, so findings are highly site specific limiting generalisation. However, this study may be useful as a model for intervention planning elsewhere, and the data is available for meta-analyses. One strength was deerstalker involvement, whose submissions enabled responsive site selection and contributed to SDNP wide tick hazard mapping. Another was our use of Density of Ticks, rather than only Density of Nymphs, which starting in the USA with Lyme disease is more common. *H.punctata*presence illustrates its reduced appropriateness for European work. Transovarial transmission of spotted fever group rickettsiae is established, so a hazard metric that excludes larvae (as Density of Nymphs does) will under-count vector density for spotted fever group rickettsiae and other pathogens (e.g. *B.miyamotoi* (Hansford *et al*., 2015)). Similarly, tick-borne encephalitis virus cycles require larvae-nymph co-feeding (PHE, 2019). The idea, based on US data, that transovarial transmission of *B.burgdorferi* s.l. is rare or non-existent (Ostfeld, 2011, p.43) is the main basis on which Density of Nymphs has been used for Lyme disease site hazard assessment. However, van Duijvendijk *et al*. (2016) showed a *Borellia* commonly causing Lyme disease in Europe can be transmitted by larvae, and in a UK study by Hall *et al*. (2017), 0.7% were *B.burgdorferi* s.l. positive. A study weakness concerns number and months of repeat sampling. Firstly, our first four sites were intended to be drag-sampled twice per year, for two years. However, this was not possible for one site, which was sampled three times only. Secondly, some species lifecycles cause seasonal differences in questing tick numbers/life-stage proportions. However, for logistical reasons the months’ individual site samplings took place in varied. This reduces confidence in validity of site comparisons somewhat, though this is partially balanced by in-year and multiple-year repeat samplings. GIS tick hazard mapping successfully brought together all publicly available, and some unpublished, tick records from SDNP. However, to our knowledge vouchers are unavailable for the historic and pan-species records included, leaving some uncertainty regarding correct species identification. In contrast a strength of our fieldwork is our vouchers.

### Conclusions

We set out to map tick hazard distribution across the SDNP, analyse habitat associations, identify and describe potential key locations for future interventions and determine their tick hazard (species and density). Against a background of increased concern about tick-borne pathogens in the UK (*B.burgdorferi* s.l., *B.miyamotoi*, *B.venatorum,* tick-borne encephalitis virus, spotted fever group rickettsiae), our mapping shows tick hazard is broadly distributed across SDNP.

*Ixodes ricinus* was the most common tick found, though the potential range expansion of *H.punctata* from its historic foci at SDNPs far east is concerning, not least as it seems better able to thrive on grazed downland than *I.ricinus*. Our study confirms woodland is the habitat in the SDNP most associated with tick hazard, but ticks (including *I.ricinus*) were collected on downland, and if *H.punctata* is allowed to expand its range westward, this is only likely to increase. Tick hazard does not reflect negatively on land managers, but should be a stimulus for action, especially at those sites with high tick risk: i.e. those with high DOT/DON and high visitor numbers. We identified key potential sites for interventions and based on measured tick-density, site description, and visitor levels have provided site specific recommendations for control measures (which should be evaluated in-situ during roll-out) and future research. These include targeted management at small high tick hazard plots with heavy visitor numbers (Queen Elizabeth Country Park), signage to increase awareness of post-visit precautions (all sites), repellent impregnated clothing for deerstalkers (Cowdray Estate), and flock-based experimental trials to control *H.punctata* (Seven Sisters Country Park, Ditchling Beacon Nature Reserve).

Further research at one of the sites with very high tick density (The Mens) may valuably contribute to an understanding of ecological dynamics underlying infection density, and potential use of predator re-introduction and protection as a public health intervention. Ecological research on *H.punctata* would also contribute towards control strategies. Whilst interventions are necessarily site-specific, this does create the danger of implementation becoming fragmented. However, SDNPA is ideally placed to link and champion site-based and regional policies to reduce hazard, whilst avoiding or reducing conflict between public health and ecosystem health.

## Acknowledgements

We thank the British Deer Society deerstalkers who submitted ticks, and organisations that enabled access to land they manage and associated data such as on visitor numbers (East Sussex County Council, Hampshire County Council, Sussex Wildlife Trust, and Cowdray Estate) and reserve pan-species records (Sussex Wildlife Trust). We are grateful to the British Deer Society grant board for methodological input, for lab support provided by technicians at Pharmacy and Biomolecular Sciences, University of Brighton, and to Jessica Stockdale who created *Fig. 7*. Finally, JM is thankful for invaluable taxonomic training and guidance provided by Professor Hans Klompen and team at Ohio State University Acarology Lab.

## Authors contributions

Author contributions detailed using CRediT Contributor Taxonomy (casrai.org/CRediT). Conceptualization, and funding acquisition: JM. Data curation: JM. Formal analysis: JM. Investigation: JM. Methodology: JM, IC, ASR. Supervision: IC, ASR. Writing – original draft: JM. Visualization: JM, ASR. Writing – review & editing: JM, IC, ASR.

## Ethical approval

Not required.

