## Supplementary Material for "Tick hazard in the South Downs National Park (UK): species, distribution, key locations for future interventions, site density, habitats"

| **Contents** |  |
| --- | --- |
| Safety measures | 2 |
| GIS layer generation  Justification of analysis | 3  3 |
| Table S1: Ticks collected at Seven Sisters Country Park, 2015 and 2016. | 4 |
| Table S2: Ticks collected at Queen Elizabeth Country Park, 2015 and 2016. | 5 |
| Table S3: Ticks collected at The Mens, 2015 and 2016. | 6 |
| Table S4: Ticks collected at Cowdray Estate, 2015 and 2016. | 7 |
| Figure S1: Magnitudes and directions of differences from post-hoc Dunn's Test of nymph and tick (all life stages) hazard at four sites sampled both 2015 and 2016. | 8 |
| Figure S2: Magnitudes and directions of differences from a post-hoc Dunn's Test in tick (all life stages) hazard at transects (n=24) coded to habitat sampled both 2015 and 2016. | 8 |

**Safety measures**

*Researcher safety and cross-contamination control*

So JM could better see and remove any ticks that attached to them during drag-sampling, white collection clothes and wellington boots (taped to trousers) were worn. These were removed on-site, and sealed in a container with the collection materials. Clothing and collection blankets, chaps, and flags were machine washed at high heat and dried between uses, which reliably kills ticks (Nelson *et al.*, 2016). These measures also reduced the risk of cross-contaminating sites with tick or pathogen species, a potential hazard parallel to concerns raised by Dunn (2014) about invasive aquatic species and inadequate biosecurity in ecological fieldwork.

JM was not aware of having been bitten during the study, or developing any signs or symptoms indicating borreliosis. Blood samples were taken at JM’s general practice before field season one, between field seasons, and after field season two. LD seropositivity was tested for at Brighton and Sussex University Hospitals NHS Trust. All tests were negative, suggesting JM had not contracted borreliosis during the study.

Where transects chosen through randomisation would have been dangerous to drag-sample randomisation was repeated until OS Grid references for safe transects arose. This was required once at Ditchling Beacon (very steep slope over a road) and twice at Seven Sisters Country Park (in the waters of Cuckmere).

Dunn A. 2014 Invasive species and invasive diseases: parallels and interactions. Disease Ecology: British Society of Parisitology Autumn Symposium 2014, 18 September 2017 University of Salford.

Nelson CA, Hayes CM, Markowitz MA, Flynn JJ, Graham AC, Delorey MJ, Mead PS, Dolan MC. 2016. The heat is on: Killing blacklegged ticks in residential washers and dryers to prevent tickborne diseases. Ticks and Tick-borne diseases 7:958. DOI: 10.1016/j.ttbdis.2016.04.016.

*Deerstalker submissions*

Deerstalkers were asked to put on disposable gloves, inspect the whole animal (stating ‘part exam’ on the form if that was not possible), collect every visible tick (using the supplied ‘tick twister’), and place them in pre-coded 1ml cryovials, pre-filled with 70% ethanol, before disposing of gloves and washing hands/using supplied alcohol hand gel. Deerstalkers were given information sheets from Public Health England on signs and symptoms to look out for and when to seek general practitioner advice.

Cryovals containing ticks were returned by post in pre-prepared, pre-paid envelopes within sealed bags, wrapped in bubble wrap and absorbent secondary packaging (in line with Royal Mail requirments).

**GIS layer generation**

In addition to individual ticks submitted by the public, Public Health England/Health Protection Agency maps also include (marked as such) the historic biodiversity records now also available in National Biodiversity Network Atlas. However, Public Health England/Health Protection Agency maps are at 10 km^2^ resolution, which does not allow confirmation records are from within the SDNP for those 10 km grid squares at its border which include land outside as well as in the Park. Given this, historic data was taken from NBN Atlas which has point records. Where historic point records were outside the Park, grid squares involved were not marked positively for recorded presence. The same approach was taken to point data extracted from Medlock et al. (2018). Those individual OS 10 km grid squares with presence records from pre-2005 only were highlighted on our maps with latest record dates.

**Justification of analysis**

Non-parametric testing was carried out to determine the significance of any difference between tick hazards (1) at the four sites, and (2) at transects with differing coded habitat types. Non-parametric testing was used for two reasons. Firstly, the number of samples covering the four sites (90 samplings, 24 transects) may not have adequately captured a normal population distribution, if one was present. Secondly, given the study concerned the collection of parasitic organisms (often grouped) whose exact siting is highly dependent on placement by sparse hosts, data would not be expected to be normally distributed. Ditchling Beacon Nature Reserve, sampled only in 2016, was not included in statistical analysis as conclusions based on single year tick samplings have limited validity (Dobson*,* Taylor & Randolph, 2011).

Dobson ADM, Taylor JL, Randolph SE. 2011. Tick (Ixodes ricinus) abundance and seasonality at recreational sites in the UK: Hazards in relation to fine-scale habitat types revealed by complementary sampling methods. Ticks and Tick-borne Diseases 2:67–74. DOI: 10.1016/j.ttbdis.2011.03.002.

**Table S1:**

**Ticks collected at Seven Sisters Country Park, 2015 and 2016.**

Ticks were collected through combined sampling with woollen blanket (B), chap (C), and flags (F). NC=not collected.

| Plot  and  Habitat | Dominant vegetation; under-growth;  main litter | Date | Undergrowth (height cm) | Rh % | | T ^O^C | | Ticks Collected  (*H.punctata*) | | | |
| --- | --- | --- | --- | --- | --- | --- | --- | --- | --- | --- | --- |
|  |  |  |  | **50cm** | **Litter** | **50cm** | **Litter** | **Larvae** | **Nymphs** | **Adults** | **Totals** |
| *Fig.* 1A  Downland  (sheep grazed, busy footpath) | Grass; dense grass; no visible litter | 5.6.15 | 70 | 69 | 74 | 20 | 26 |  |  |  | 0 |
|  |  | 29.9.15 | 10 | 57 | 90 | 17 | 20 |  |  |  | 0 |
|  |  | 2015 transect totals | | | | | |  |  |  | 0 |
|  |  | 30.4.16 | 5 | 60 | 74 | 11 | 11 |  |  |  | 0 |
|  |  | 24.8.16 | NC | 43 | 50 | 29 | 28 |  |  |  | 0 |
|  |  | 2016 transect totals | | | | | |  |  |  | 0 |
|  |  | **Transect totals (range)** | | | | | |  |  |  | **0 (0-0)** |
| *Fig*. 1B  Downland  (sheep grazed) | Grass; dense grasses, nettles; no visible litter | 5.6.15 | 0 | 75 | 80 | 19 | 19 |  |  |  | 0 |
|  |  | 29.9.15 | 3-5 | 57 | 67 | 18 | 19 |  |  |  | 0 |
|  |  | 2015 transect totals | | | | | |  |  |  | 0 |
|  |  | 30.4.16 | 2-15 | 62 | 65 | 10 | 12 |  |  |  | 0 |
|  |  | 24.8.16 | NC | 42 | 62 | 30 | 31 |  |  |  | 0 |
|  |  | 2016 transect totals | | | | | |  |  |  | 0 |
|  |  | **Transect totals (range)** | | | | | |  |  |  | **0 (0-0)** |
| *Fig*. 1C  Downland  (sheep grazed) | Grass; dense grass; no visible litter | 5.6.15 | 0 | 66 | 76 | 17 | 19 |  | 2B |  | 2B |
|  |  | 29.9.15 | 15-30 | 57 | 66 | 17 | 16 |  |  |  | 0 |
|  |  | 2015 transect totals | | | | | |  | 2 |  | 2 |
|  |  | 30.4.16 | 5 | 52 | 83 | 11 | 11 |  |  |  | 0 |
|  |  | 24.8.16 | 35 | 45 | 50 | 29 | 31 |  |  |  | 0 |
|  |  | 2016 transect totals | | | | | |  |  |  | 0 |
|  |  | **Transect totals (range)** | | | | | |  | **2** |  | **2 (0-2)** |
| *Fig*. 1D  Woodland  (High canopy, carpark picnic area) | Beech, conifer; no undergrowth; conifer and beach litter | 10.6.15 | 0 | 50 | 52 | 17 | 17 |  |  |  | 0 |
|  |  | 29.9.15 | 0 | 57 | 75 | 17 | 16 |  |  |  | 0 |
|  |  | 2015 transect totals | | | | | |  |  |  | 0 |
|  |  | 30.4.16 | 0 | 49 | 83 | 12 | 12 |  |  |  | 0 |
|  |  | 24.8.16 | 0 | 49 | 56 | 26 | 26 |  |  |  | 0 |
|  |  | 2016 transect totals | | | | | |  |  |  | 0 |
|  |  | **Transect totals (range)** | | | | | |  |  |  | **0 (0-0)** |
| *Fig*. 1E  Woodland  (multi-level canopy, next to visitor carpark) | Sycamore; sycamore saplings, nettles, ivy; sycamore leaves | 10.6.15 | 100 | 46 | 50 | 18 | 18 |  |  |  | 0 |
|  |  | 29.9.15 | 100 | 62 | 80 | 17 | 16 |  |  |  | 0 |
|  |  | 2015 transect totals | | | | | |  |  |  | 0 |
|  |  | 30.4.16 | 15-100 | 50 | 69 | 12 | 14 |  |  |  | 0 |
|  |  | 24.8.16 | 100 | 52 | 56 | 27 | 25 |  |  |  | 0 |
|  |  | 2016 transect totals | | | | | |  |  |  | 0 |
|  |  | **Transect totals (range)** | | | | | |  |  |  | **0 (0-0)** |
| *Fig*. 1F  Downland  (sheep grazed, busy path) | Grass; thistles, dense grasses; no visible litter | 5.6.15 | 50 | 56 | 67 | 18 | 20 |  |  | 2♀C | 2C |
|  |  | 29.9.15 | 50 | 59 | 87 | 19 | 20 |  |  |  | 0 |
|  |  | 2015 transect totals | | | | | |  |  | 2 | 2 |
|  |  | 30.4.16 | 5 | 44 | 65 | 12 | 14 |  |  |  | 0 |
|  |  | 24.8.16 | 6 | 35 | 42 | 31 | 32 |  |  |  | 0 |
|  |  | 2016 transect totals | | | | | |  |  |  | 0 |
|  |  | **Transect totals (range)** | | | | | |  |  | **2** | **2(0-2)** |
| Site transect totals  (mean, range, IQR) | | | | | | | |  | **2** | **2♀** | **4 (1,**  **0-2, 0-2)** |
| Extras | | 2015 (all B) | | | | | |  | 6 | 1♀ | 7 |
|  |  | 2016 (all B) | | | | | | 65 | 2 |  | 67 |
|  |  | **Total extra ticks collected** | | | | | | **65** | **8** | **1♀** | **74** |
| Total ticks collected at site | | | | | | | | **65** | **10** | **3♀** | **78** |

**Table S2:**

**Ticks collected at Queen Elizabeth Country Park, 2015 and 2016.**

Ticks were collected through combined sampling with woollen blanket (B), chap (C), and flags (F). *2 removed for safety validation before light microscopy confirmation of ID. NC=not collected. SDW = South Downs Way national trail.

| Plot  and  Habitat | Dominant vegetation; under-growth;  main litter | Date | Undergrowth (height cm) | Rh % | | T ^O^C | | Ticks Collected  (*I.ricinus*) | | | |
| --- | --- | --- | --- | --- | --- | --- | --- | --- | --- | --- | --- |
|  |  |  |  | **50cm** | **Litter** | **50cm** | **Litter** | **Larvae** | **Nymphs** | **Adults** | **Totals** |
| *Fig*. 1G  Woodland  (high canopy, SDW from visitor cnt.) | Beech, ash; nettles; beech litter | 2.8.15 | 40 | 39 | 44 | 24 | 24 |  | 1B |  | 1 |
|  |  | 6.9.15 | 80 | 54 | 69 | 14 | 15 |  | 1B | 1♀B | 2B |
|  |  | 2015 transect totals | | | | | |  | 2 | 1 | 3 |
|  |  | 24.4.16 | nc | 56 | 82 | 9 | 9 |  | 2B |  | 2B |
|  |  | 24.9.16 | nc | 67 | 83 | 18 | 17 |  | 3B |  | 3B |
|  |  | 2016 transect totals | | | | | |  | 5 |  | 5 |
|  |  | **Transect totals (range)** | | | | | |  | **7** | **1** | **8 (0-3)** |
| *Fig.* 1H  Woodland  (high canopy) | Conifer, beech, yew; nettle patches; conifer & beach litter | 2.8.15 | 0 | 44 | 55 | 22 | 22 |  |  |  | 0 |
|  |  | 6.9.15 | 0-80 | 59 | 76 | 15 | NC | 2B | 1B |  | 3B |
|  |  | 2015 transect totals | | | | | | 2 | 1 |  | 3 |
|  |  | 24.4.16 | 0-50 | 49 | 81 | 10 | 9 |  |  |  | 0 |
|  |  | 24.9.16 | 0-100 | 66 | 99 | 18 | 16 | 10B | 3B |  | 13B |
|  |  | 2016 transect totals | | | | | | 10 | 3 |  | 13 |
|  |  | **Transect totals (range)** | | | | | | **12** | **4** |  | **16 (0-13)** |
| *Fig*. 1I  Woodland  (high canopy) | Beech, yew, conifer; brambles, grass; beech and conifer litter | 15.6.15 | 0-20 | 61 | 69 | 17 | 16 |  | 1B, 1C |  | 1B, 1C |
|  |  | 2.8.15 | 60 | 50 | 71 | 22 | 24 |  | 1B, 1C | 1♂B | 2B, 1C |
|  |  | 2015 transect totals | | | | | |  | 4 | 1 | 5 |
|  |  | 24.4.16 | 60 | 45 | 83 | 9 | 11 |  | 8B, 2C | 1♂B | 9B, 2C |
|  |  | 24.9.16 | nc | 68 | 88 | 19 | 17 | 24B | 1B |  | 25B |
|  |  | 2016 transect totals | | | | | | 24 | 11 | 1 | 36 |
|  |  | **Transect totals (range)** | | | | | | **24** | **15** | **2** | **41 (2-25)** |
| *Fig.* 1J  Woodland  (high canopy, 20m from picnic area) | Beech; beech saplings, nettles; beach mast and leaves, | 15.6.15 | 0-100 | 57 | 66 | 18 | 17 |  | 1B |  | 1B |
|  |  | 2.8.15 | 30 | 46 | 72 | 20 | 20 |  |  |  | 0 |
|  |  | 2015 transect totals | | | | | |  | 1 |  | 1 |
|  |  | 24.4.16 | 0 | 39 | 52 | 11 | 12 |  |  |  | 0 |
|  |  | 24.9.16 | 0 | 60 | 84 | 19 | 17 |  |  |  | 0 |
|  |  | 2016 transect totals | | | | | |  |  |  | 0 |
|  |  | **Transect totals (range)** | | | | | |  | **1** |  | **1 (0-1)** |
| *Fig.* 1K  Woodland  (footpath from car park) | Grass and nettles; | 15.6.15 | 70 | 66 | 72 | 18 | 17 |  |  |  | 0 |
|  |  | 2.8.15 | 100 | 43 | 53 | 21 | 21 |  |  |  | 0 |
|  |  | 2015 transect totals | | | | | |  |  |  | 0 |
|  |  | 24.4.16 | 21 | 45 | 86 | 10 | 11 |  |  |  | 0 |
|  |  | 24.9.16 | 10-75 | 71 | 86 | 18 | 18 |  |  |  | 0 |
|  |  | 2016 transect totals | | | | | |  |  |  | 0 |
|  |  | **Transect totals (range)** | | | | | |  |  |  | **0 (0-0)** |
| *Fig*. 1L  Woodland  (low canopy) | Beech, hazel, yew; brambles, beech saplings; beech litter | 2.8.15 | 20 | 54 | 60 | 20 | 20 |  | 5B |  | 5B |
|  |  | 6.9.15 | 20 | 63 | 73 | 14 | 14 |  |  |  | 0 |
|  |  | 2015 transect totals | | | | | |  | 5 |  | 5 |
|  |  | 24.4.16 | 60 | 59 | 90 | 9 | 9 |  | 10B, 4C | 1♂B | 11B, 4C |
|  |  | 24.9.16 | nc | 79 | 88 | 17 | 16 | 1B | 1B | 1♂B | 3B |
|  |  | 2016 transect totals | | | | | | 1 | 15 | 2 | 18 |
|  |  | **Transect totals (range)** | | | | | | **1** | **20** | **2** | **23 (1-15)** |
| Site transect totals  (mean, range, IQR) | | | | | | | | **37** | **47** | **1♀, 4♂** | **89 (15,**  **0-41,1-28)** |
| Extras | | 2015 (all B) | | | | | | 16 | 45* | 2♀ | 63 |
|  |  | 2016 (all B) | | | | | | 9 | 20* | 1♀, 1♂ | 31 |
|  |  | **Total extra ticks collected** | | | | | | **25** | **65** | **3♀, 1♂** | **94** |
| Total ticks collected at site | | | | | | | | **62** | **112** | **4♀, 5♂** | **183** |

**Table S3:**

**Ticks collected at The Mens, 2015 and 2016.**

Ticks were collected through combined sampling with woollen blanket (B), chap (C), and flags (F). *2 removed for safety validation before light microscopy confirmation of ID. NC=not collected.

| Plot  and  Habitat | Dominant vegetation; under-growth;  main litter | Date | Undergrowth (height cm) | Rh % | | T ^O^C | | Ticks Collected  (*I.ricinus*) | | | |
| --- | --- | --- | --- | --- | --- | --- | --- | --- | --- | --- | --- |
|  |  |  |  | **50cm** | **Litter** | **50cm** | **Litter** | **Larvae** | **Nymphs** | **Adults** | **Totals** |
| *Fig*. 2A  Woodland  (footpath from car park with grass verge) | Beech; grass verge on path; beech mast and leaves | 10.5.15 | 30 | 67 | 72 | 17 | 16 |  | 10B | 2♂B | 12B |
|  |  | 19.6.15 | 0 | 59 | 63 | 16 | 16 |  | 2B |  | 2B |
|  |  | 2015 transect totals | | | | | |  | 12 | 2 | 14B |
|  |  | 23.4.16 | 5-37 | 54 | 74 | 10 | 10 |  | 12B |  | 12B |
|  |  | 30.8.16 | 15 | 66 | 88 | 19 | 18 | 1B | 1B |  | 2B |
|  |  | 2016 transect totals | | | | | | 1 | 13 |  | 14B |
|  |  | **Transect totals (range)** | | | | | | **1** | **25** | **2** | **28 (2-12)** |
| *Fig.* 2B  Woodland  (high canopy) | Beech; sparse holly in vicinity, not on plot;  beech mast and leaves, holly leaves | 10.5.15 | 0 | 68 | 76 | 16 | 17 |  | 3B |  | 3B |
|  |  | 19.6.15 | 0 | 59 | 63 | 16 | 16 |  | 1B |  | 1B |
|  |  | 2015 transect totals | | | | | |  | 4 |  | 4 |
|  |  | 23.4.16 | 0 | 41 | 84 | 10 | 11 |  |  |  | 0 |
|  |  | 30.8.16 | 0 | 65 | 84 | 20 | 19 | 4B | 2B |  | 6B |
|  |  | 2016 transect totals | | | | | | 4 | 2 |  | 6 |
|  |  | **Transect totals (range)** | | | | | | **4** | **6** |  | **10 (0-6)** |
| *Fig*. 2C  Woodland  (multi-level canopy) | Beech, oak; holly samplings; beach and oak mast and leaves, holly leaves | 10.5.15 | 50-60 | 60 | 76 | 17 | 17 |  | 1C,7B | 1♂B | 8B, 1C |
|  |  | 19.6.15 | nc | 58 | 61 | 17 | 17 |  | 1C,1B |  | 1B, 1C |
|  |  | 2015 transect totals | | | | | |  | 10 | 1 | 11 |
|  |  | 23.4.16 | 20-50 | 33 | 78 | 13 | 13 |  | 3B |  | 3B |
|  |  | 30.8.16 | 20 | 69 | 80 | 20 | 19 | 6B | 4B |  | 10B |
|  |  | 2016 transect totals | | | | | | 6 | 7 |  | 13 |
|  |  | **Transect totals (range)** | | | | | | **6** | **17** | **1** | **24 (1-10)** |
| *Fig*. 2D  Woodland  (low canopy) | Oak, beech, ash; grass, bluebells, brambles; dense grass | 10.5.15 | nc | 60 | 68 | 17 | 16 |  | 2B |  | 2B |
|  |  | 19.6.15 | 75 | 49 | 50 | 18 | 19 |  | 3B |  | 3B |
|  |  | 2015 transect totals | | | | | |  | 5 |  | 5 |
|  |  | 23.4.16 | 24-87 | 38 | 80 | 11 | 10 | 3B | 2C, 4B |  | 2C, 7B |
|  |  | 30.8.16 | 80 | 76 | 82 | nc | 20 |  |  |  | 0 |
|  |  | 2016 transect totals | | | | | | 3 | 6 |  | 9 |
|  |  | **Transect totals (range)** | | | | | | **3** | **11** |  | **14 (0-9)** |
| *Fig*. 2E  Woodland  (footpath) | Beech, oak; hawthorn, holly; dense grass | 10.5.15 | nc | 61 | 61 | 16 | 16 |  | 2B |  | 2B |
|  |  | 19.6.15 | 30 | 57 | 62 | 19 | 19 |  |  |  | 0 |
|  |  | 2015 transect totals | | | | | |  | 2 |  |  |
|  |  | 23.4.16 | 28 | 47 | 82 | 10 | 10 | 1B | 3B | 1♀B | 5B |
|  |  | 30.8.16 | 15 | 74 | 77 | 19 | 19 | 1B | 2B |  | 3B |
|  |  | 2016 transect totals | | | | | | 2 | 5 | 1 | 8 |
|  |  | **Transect totals (range)** | | | | | | **2** | **7** | **1** | **10 (0-5)** |
| *Fig*. 2F  Woodland  (multi-level canopy) | Beech, oak; holly, ferns; beach and oak mast and leaves | 10.5.16 | 0 | 66 | 74 | 16 | 16 |  | 4B* |  | 4B |
|  |  | 19.6.15 | 30 | 55 | 59 | 19 | 19 |  | 2B |  | 2B |
|  |  | 2015 transect totals | | | | | |  | 6 |  | 6 |
|  |  | 23.4.16 | 20 | 41 | 75 | 10 | 9 |  | 8B |  | 8B |
|  |  | 30.8.16 | 10-30 | 74 | 80 | 20 | 20 | 19B | 2B |  | 21B |
|  |  | 2016 transect totals | | | | | | 19 | 10 |  | 29 |
|  |  | **Transect totals (range)** | | | | | | **19** | **16** |  | **35 (2-21)** |
| Transect totals  (range, IQR) | | | | | | | | **35** | **82** | **1♀, 3♂** | **121 (10-35,**  **10-30)** |
| Extras | | 2015 (all B) | | | | | | 29 | 54 | 1♀, 3♂ | 87 |
|  |  | 2016 (all B) | | | | | | 71 | 50 | 1♀ | 122 |
|  |  | **Total extra ticks collected** | | | | | | **100** | **104** | **2♀, 3♂** | **209** |
| Total ticks collected at site | | | | | | | | **135** | **186** | **3♀, 6♂** | **330** |

**Table S4:**

**Ticks collected at Cowdray Estate, 2015 and 2016.**

Ticks were collected through combined sampling with woollen blanket (B), chap (C), and flags (F).

*1 removed for safety validation before light microscopy confirmation of ID. NC=not collected. SDW = South Downs Way national trail.

| Plot  and  Habitat | Dominant vegetation; under-growth;  main litter | Date | Undergrowth (height cm) | Rh % | | T ^O^C | | Ticks Collected  (*I.ricinus*) | | | |
| --- | --- | --- | --- | --- | --- | --- | --- | --- | --- | --- | --- |
|  |  |  |  | **50cm** | **Litter** | **50cm** | **Litter** | **Larvae** | **Nymphs** | **Adults** | **Totals** |
| *Fig*. 2G  Conifer-Heath | Conifers (recently planted); dense grass; no visible litter | 15. 5.15 | 50 | 55 | 78 | 18 | 23 |  | 2B*,  1C |  | 2B, 1C |
|  |  | 15.6.15 | 75 | 66 | 72 | 15 | nc |  |  |  | 0 |
|  |  | 2015 transect totals | | | | | |  | 3 |  | 3 |
|  |  | 15.11.16 | 100-200 | 56 | 82 | 17 | 16 |  |  |  | 0 |
|  |  | 2016 transect totals | | | | | |  |  |  | 0 |
|  |  | **Transect totals (range)** | | | | | |  | **3** |  | **3 (0-3)** |
| *Fig*. 2H  Conifer-heath border  (track) | Ferns, brambles; dense grass; fern leaves | 15.5.15 | 30 | 60 | 70 | 15 | 15 |  | 7B |  | 7B |
|  |  | 15.6.15 | nc | nc | nc | nc | nc |  | 1B |  | 1B |
|  |  | 2015 transect totals | | | | | |  | 8 |  | 8 |
|  |  | 15.11.16 | 10-120 | 55 | 83 | 16 | 16 |  |  |  | 0 |
|  |  | 2016 transect totals | | | | | |  |  |  | 0 |
|  |  | **Transect totals (range)** | | | | | |  | **8** |  | **8 (0-7)** |
| *Fig*. 2I  Woodland  (very dense canopy) | Conifers; no undergrowth; pine needles and branches | 15.5.15 | 0 | nc | nc | nc | Nc |  | 5B |  | 5B |
|  |  | 15.6.15 | 0 | 80 | 81 | 14 | 14 |  | 4B |  | 4B |
|  |  | 2015 transect totals | | | | | |  | 9 |  | 9 |
|  |  | 15.11.16 | 0-50 | 59 | 82 | 59 | 14 |  | 1B |  | 1B |
|  |  | 2016 transect totals | | | | | |  | 1 |  | 1 |
|  |  | **Transect totals (range)** | | | | | |  | **10** |  | **10 (1-5)** |
| *Fig*. 2J  Downland  (field bordering wood) | Grass (sheep grazed); dense grass; no visible litter | 15.5.15 | 50 | 57 | 73 | 14 | 14 |  |  | 1♂B | 1B |
|  |  | 15.6.15 | nc | 52 | 81 | 19 | 17 |  |  |  | 0 |
|  |  | 2015 transect totals | | | | | |  |  | 1 | 1 |
|  |  | 25.9.16 | 10 | 62 | 85 | 17 | 14 |  |  |  | 0 |
|  |  | 2016 transect totals | | | | | |  |  |  | 0 |
|  |  | **Transect totals (range)** | | | | | |  |  | **1** | **1 (0-1)** |
| *Fig*. 2K  Downland  (field bordering wood) | Grass (sheep grazed); dense grass; no visible litter | 15.5.15 | 30 | 63 | 76 | 15 | 16 |  |  |  | 0 |
|  |  | 15.6.15 | nc | 69 | 80 | 15 | 16 |  |  |  | 0 |
|  |  | 2015 transect totals | | | | | |  |  |  | 0 |
|  |  | 25.9.16 | 10 | 56 | 89 | 17 | 16 |  |  |  | 0 |
|  |  | 2016 transect totals | | | | | |  |  |  | 0 |
|  |  | **Transect totals (range)** | | | | | |  |  |  | **0 (0-0)** |
| *Fig*. 2L  Downland  (SDW footpath) | Nettles; dense grass; no visible litter | 15.5.15 | nc | 78 | 73 | 14 | 12 |  |  |  | 0 |
|  |  | 15.6.15 | 120 | 66 | 76 | 18 | 20 |  |  |  | 0 |
|  |  | 2015 transect totals | | | | | |  |  |  | 0 |
|  |  | 25.9.16 | 100 | 59 | 90 | 16 | 16 |  | 1B |  | 1B |
|  |  | 2016 transect totals | | | | | |  | 1 |  | 1 |
|  |  | **Transect totals (range)** | | | | | |  | **1** |  | **1 (0-1)** |
| Site transect totals  (mean, range, IQR) [Three samplings only, due to a 2016 access restriction] | | | | | | | |  | **22** | **1♂** | **23 (4,**  **0-10, 1-9)** |
| Extras | | 2015 (all B) | | | | | |  | 2 |  | 2 |
|  |  | 2016 (all B) | | | | | | 3 | 3 |  | 6 |
|  |  | **Total extra ticks collected** | | | | | | **3** | **5** |  | **8** |
| Total ticks collected at site | | | | | | | | **3** | **27** | **1♂** | **31** |

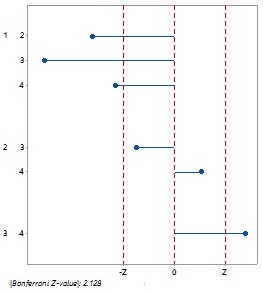

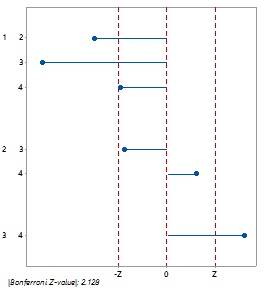

**Figure S1:**

**Magnitudes and directions of differences from post-hoc Dunn's Tests on nymph and tick (all life stages) hazard at four sites sampled both 2015 and 2016.**

A) Pairwise comparison of site nymph hazard. (B) Pairwise comparison of site tick (all life stages). 1=The Mens; 2=Cowdray Estate; 3=Seven Sisters Country Park; 4=Queen Elizabeth Country Park.

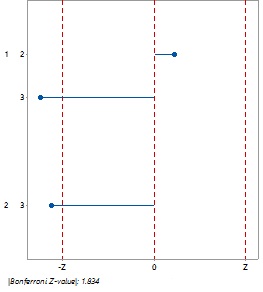

**Figure S2:**

**Magnitudes and directions of differences from a post-hoc Dunn's Test on tick (all life stages) hazard at transects (n=24) coded to habitat sampled both 2015 and 2016.**

1=deciduous woodland; 2=Conifer woodland/planting; 3=Sheep grazed downland.
